# Oxytocin and Dopamine Receptor Expression: Cellular Level Implications for Pair Bonding

**DOI:** 10.1101/2025.03.03.640889

**Authors:** Meredith K. Loth, Julia C. Schmidt, Cassandra A. Gonzalez, Liza E. Brusman, Julie M. Sadino, Kelly E. Winther, David S.W. Protter, Zoe R. Donaldson

**Author notes:** corresponding author: Zoe R. Donaldson, PhD, University of Colorado Boulder, 347 UCB, Boulder, CO 80309, USA.

## Abstract

Oxytocin (*Oxtr*) and dopamine (*Drd1*, *Drd2*) receptors provide a canonical example for how differences in neuromodulatory receptors drive individual and species-level behavioral variation. These systems exhibit striking and functionally-relevant differences in nucleus accumbens (NAc) expression across monogamous prairie voles (*Microtus ochrogaster*) and promiscuous meadow voles (*Microtus pennsylvanicus*). However, their cellular organization remains largely unknown. Using multiplex *in situ* hybridization, we mapped *Oxtr*, *Drd1*, and *Drd2* expression in sexually naïve and mate-paired prairie and meadow voles. Prairie voles have more *Oxtr+* cells than meadow voles, but *Oxtr* distribution across dopamine-receptor cell class was similar, indicating a general upregulation rather than cell class bias. *Oxtr* was enriched in cells that express both dopamine receptors (*Drd1+/Drd2+*) in prairie voles, suggesting these cells may be particularly sensitive to oxytocin. We found no species or pairing-induced differences in *Drd1+* or *Drd2+* cell counts, suggesting prior reports of expression differences may reflect upregulation in cells already expressing these receptors. Finally, we used single-nucleus sequencing to provide the first comprehensive map of *Oxtr* and *Drd1-5* across molecularly-defined NAc cell types in the prairie vole. These results provide a critical framework for understanding how nonapeptide and catecholamine systems may recruit distinct NAc cell types to shape social behavior.

## 1. Introduction

Oxytocin and dopamine, acting via their receptors, play critical roles in regulating complex social behaviors, including pair bonding, parental care, and social recognition[1–3]. Comparisons of oxytocin (*Oxtr* - mRNA; OXTR - protein) and dopamine (*Drd1* and *Drd2* – mRNA; DRD1 and DRD2 - protein) receptor distributions within and across species provide classic examples of how differences in receptor distribution and/or levels within the brain contribute to species-appropriate and experience-dependent behaviors such as mating and aggression[4][5]. Socially monogamous prairie voles (*Microtus ochrogaster*) and promiscuous meadow voles (*Microtus pennsylvanicus*) exemplify this relationship, exhibiting strikingly different social behavior alongside distinct limbic oxytocin and dopamine receptor expression[6–9].

Prairie voles form selective and long-lasting pair bonds, exhibit aggression toward non-partner individuals, and display robust biparental care, whereas meadow voles are primarily asocial, do not form mating-based bonds, and rely solely on dams for pup-rearing[10]. Compared to meadow voles, prairie voles have higher OXTR densities in the nucleus accumbens (NAc), which functionally contribute to species differences in social behavior[6,11–13]. Blockade of these receptors impairs bond formation, while overexpression accelerates bonding and facilitates alloparenting[12][14,15]. Additionally, histone deacetylase inhibitors facilitate partner preference formation, upregulating *Oxtr* transcript and OXTR protein, suggesting epigenetic modulation of social bonding that may be mediated via OXTR[16]. Of note, artificially increasing OXTR levels in adult meadow voles is not sufficient to convey the ability to form partner preference, suggesting it may be developmentally required for its effects on social behavior in prairie voles[17].

Dopamine receptor expression in the NAc also differs across species and social experiences. DRD1-class (DRD1+DRD5) and DRD2-class (DRD2+DRD3+DRD4) receptors play opposing roles in pair bonding in prairie voles, and their balance in the NAc is critical for modulating reward signaling and partner preference[18,19]. DRD2-class agonists promote partner preference formation, and DRD2-class antagonists block mating-induced partner preference formation[20,21]. Conversely, pharmacological manipulation of DRD1 receptors in prairie voles has opposite effects; agonism blocks pair bonding and leads to enhanced aggression[18,22]. DRD1 receptors are also upregulated following pair bond formation, potentially reinforcing social attachment by promoting aggressive rejection of non-partner voles[23]. In line with this, meadow voles exhibit higher DRD1 levels, which may contribute to their lack of bonding after mating[18]. Thus, the balance between DRD1 and DRD2—which differs across species and social experience, likely contributes to species and experience dependent differences in social bonding behaviors.

While previous studies have examined the distributions of oxytocin and dopamine receptors independently, their interaction is also critical. In prairie voles, pair bonding depends on the concurrent activation of OXTR and DRD2 in the NAc[20]. Studies to date have largely quantified receptors levels via autoradiography, or in a few cases, qRT-PCR or western blotting [2,6,8,9,13,24]. These and initial papers examining mRNA via radiolabeled *in situ* hybridization fall short of providing cell-type specific information regarding the expression of *Oxtr, Drd1,* and *Drd2* mRNA expression. Recently, however, more sensitive techniques such as fluorescent *in situ* hybridization (FISH) has enabled higher-resolution analyses of receptor localization, revealing that *Oxtr* transcripts can be co-expressed with either *Drd1* or *Drd2* in sexually naïve prairie voles[25]. However, it remains unclear whether this co-expression is unique to prairie voles and/or if sociosexual experiences—such as pair bonding—alter its incidence or frequency. Addressing this question is critical for understanding the cellular basis of social bonding and how oxytocin-dopamine interactions support pair bonding. Furthermore, the role of these receptors in bonding is likely multifaceted, as DRD1 is not only implicated in non-partner aggression, but also in partner-directed motivation in bonded prairie voles[26]. Without a complete understanding of receptor distribution and co-expression, it remains unknown how DRD1 mediates both aggression and motivation within bonded relationships.

To explore the cellular organization of these receptors across species and social experiences, we examined *Oxtr*, *Drd1*, and *Drd2* mRNA expression using multiplex FISH. We compared sexually naïve prairie and meadow voles to their mated counterparts, a social experience that induces pair bonding in prairie but not meadow voles. Additionally, we leveraged well-powered tissue- and nuclear-level sequencing datasets to create a map that delineates the distribution of *Oxtr* and dopamine receptor transcripts (*Drd1-5*) in molecularly-defined NAc cell types in prairie voles[27]. Our analysis revealed species-specific and cell-type specific patterns of receptor expression, providing critical insights into how these neuromodulators may affect cellular and circuit function to drive differential social behaviors.

## 2. Results

All statistical results are reported in Supplementary Table 1. Only significant main effects or interactions (p < 0.05) and corresponding post-hoc tests are reported below.

### 2.1. Prairie voles have more *Oxtr+* cells than meadow voles in the NAc core and shell

A greater proportion of cells were *Oxtr+* in prairie voles compared to meadow voles across all NAc subregions. The lateral shell showed the largest species difference (34.03% prairie vs 6.81% meadow *Oxtr*+ cells). (Fig 1C: 2-way repeated measures (RM) ANOVA: main effect of transcript identity: F(_3, 72_) = 45.54, p < 0.0001; interaction [transcript identity x species]: F(_3, 72_) = 6.816, p = 0.0004. Post hoc Sidak: *Oxtr* Prairie vs *Oxtr* Meadow: p < 0.0001; Fig 1D: 2-way RM-ANOVA: main effect of transcript identity: F(_3, 72_) = 54.39, p < 0.0001; interaction [transcript identity x species]: F(_3, 69_) = 5.797, p = 0.0014). Post hoc Sidak: *Oxtr* Prairie vs *Oxtr* Meadow: p = 0.0041; Fig 1E: 2-way RM-ANOVA: main effect of transcript identity: F(_3, 72_) = 49.880, p < 0.0001; main effect of species F(_1, 24_) = 10.670, p = 0.0033; interaction [transcript identity x species]: F(_3, 72_) = 11.490, p < 0.0001). Post hoc Sidak: *Oxtr* Prairie vs *Oxtr* Meadow: p < 0.0001). We also found that a majority of cells were positive for *Drd1* or *Drd2* with no species differences in their relative proportions (Fig 1D-F). Similarly, we did not observe species differences in the proportion of cells (16 – 23%) that did not express any of the 3 labeled transcripts (Fig 1D-F).

**Figure 1.**
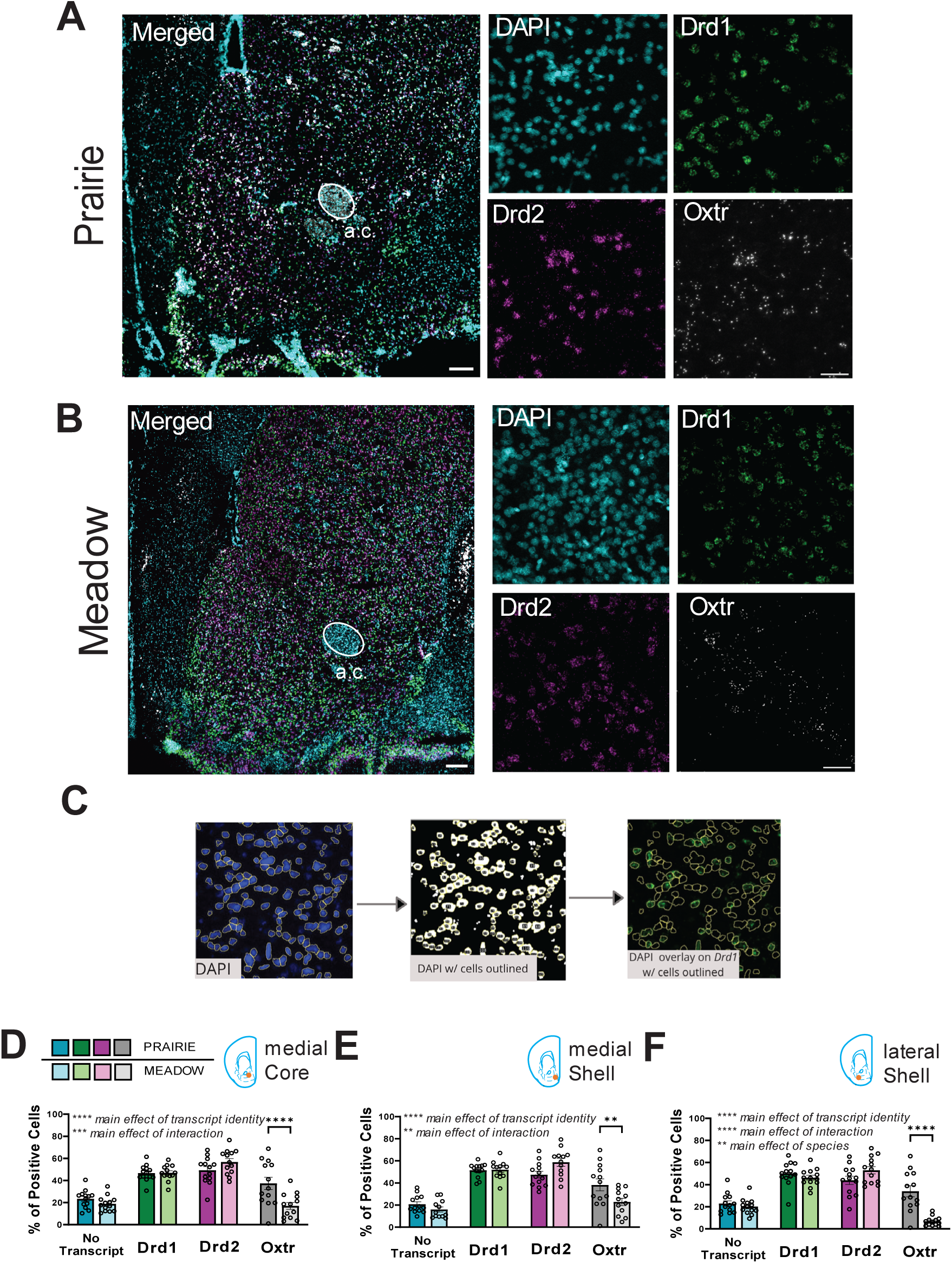
Prairie voles have more *Oxtr* positive cells than meadow voles in the nucleus accumbens. **A-B.** Representative *in situ* hybridization images of dopamine receptor 1 (*Drd1*), dopamine receptor 2 (*Drd2*), and oxytocin receptor (*Oxtr*) in the nucleus accumbens core of prairie voles (**A**) and meadow voles (**B**). Scale bar = 200 µm in low magnification images (left) and 50 µm in high magnification images (right). **C.** Cell quantification using Fiji Image J: DAPI stained cells are computationally identified and thresholding is manually adjusted to remove background and detect nuclei. Nuclei are outlined, numbered and an outline mask is generated to overlay on other channels to then identify positive cells for each transcript channel. **D-F.** Quantification of transcript presence or absence within a DAPI mask in the nucleus accumbens core (**D**), medial shell (**E**), and lateral shell (**F**) of prairie voles and meadow voles. There was a main effect of transcript identity and a significant interaction between species and transcript identity. Post-hoc tests revealed no significant differences between species for No transcript, Drd1, or Drd2 positive cells. However, prairie voles had a greater number of *Oxtr* receptor positive cells than meadow voles in all subregions examined. Error bars show SEM. n = 13 per group. ** p < 0.01, **** p < 0.0001.

We also performed *Oxtr* SNP genotyping on our prairie vole samples post-mortem (Supplementary Table 2). Across all genotyped prairie voles (n = 13), genotype distribution was 69% heterozygous, 23% high-*Oxtr*, and 8% low-*Oxtr*. Due to the presence of only one low-*Oxtr* animal, we grouped it with the heterozygous animals. Comparison of high *Oxtr* animals to non-high *Oxtr* animals revealed no statistically significant differences in receptor distribution. Therefore, we pooled all genotypes for subsequent analyses.

### 2.2. *Oxtr* co-expression with dopamine receptor cell classes is greater in prairie voles than meadow voles in the lateral shell

Next, we classified cells based on receptor expression, yielding 8 transcript combinations: No transcript; *Drd1+ only*; *Drd2+ only*; *Drd1*+*Drd2*; *Oxtr+ only*; *Drd1*+*Oxtr*; *Drd2*+*Oxtr*; and *Drd1*+*Drd2*+*Oxtr*. We replicated previous findings that *Oxtr* is expressed on both *Drd1+* and *Drd2+* cells in prairie voles (7-14 % of cells, depending on the transcript combination and subregion examined)[25] (Fig. 2A-C). Meadow voles also exhibited *Oxtr* co-expression but at significantly lower levels (2–8%, p < 0.05), consistent with lower overall *Oxtr*+ cell counts. We also found that *Oxtr* is expressed on *Drd1*+*Drd2+* cells, and this population is enriched in prairie voles in the lateral shell. Conversely, meadow voles exhibited higher proportions of *Drd2+* only and *Drd1+Drd2+* cells, while no transcript and *Drd1+* only transcript combinations did not differ significantly between species.

**Figure 2.**
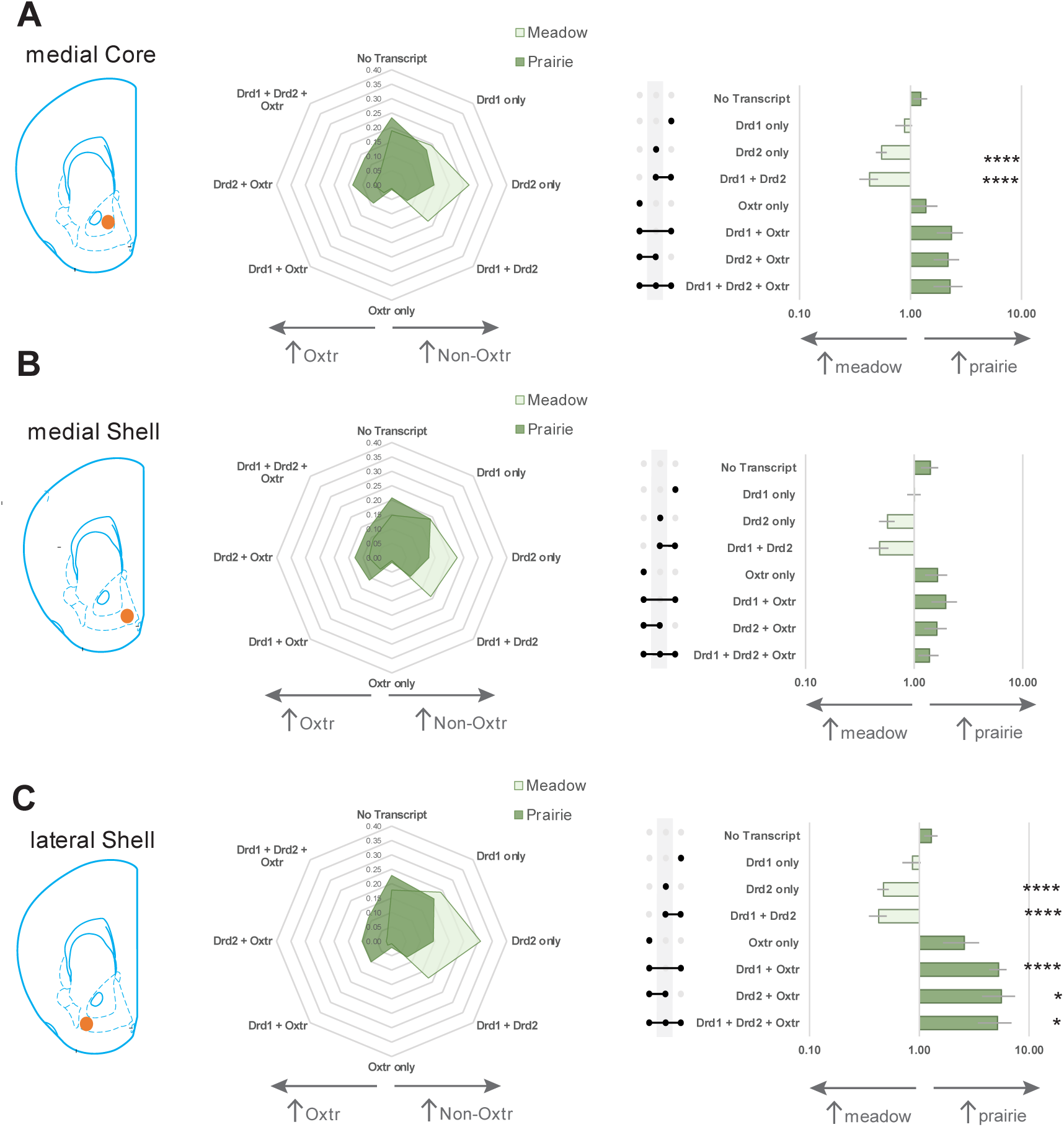
*Oxtr* co-expression with dopamine receptor cell classes is greater in prairie voles compared to meadow voles in the nucleus accumbens. **A-C.** Quantification of cell proportions based on transcript combinations the nucleus accumbens core (**A**), medial shell (**B**), and lateral shell (**C**) in meadow voles and prairie voles (light green and darker green respectively in radar plot). Fold change of prairie:meadow vole of each transcript combination was quantified and compared between species for each subregion. Meadow voles had a significantly greater number of *Drd2* only and *Drd1+Drd2* positive cells compared to prairie voles in all subregions. Prairie voles demonstrated significantly higher levels of *Oxtr* co-expressing transcripts as compared to meadow voles in the lateral shell. Co-expression of *Oxtr* does not appear to be biased towards a particular *Drd* receptor class as co-expression is found with *Drd1*, *Drd2*, and *Drd1+Drd2* cells in all subregions. Error bars show EP (error propagation). N = 12-13 per group. * p < 0.05, ** p < 0.01, *** p < 0.001, **** p < 0.0001.

To analyze differences in relative proportions of transcript combinations between species, we calculated the mean prairie:meadow vole ratio and propagated measurement errors to estimate uncertainty. Using a one-way t-test relative to null with FDR corrected p-values, we found that prairie voles have greater proportions of *Oxtr* co-expressing cells compared to meadow voles, particularly in the lateral shell (Fig 2A-C: *Drd1* + *Oxtr* Prairie vs *Drd1* + *Oxtr* Meadow: t = 4.4565, p < 0.0001; *Drd2* + *Oxtr* Prairie vs *Drd2* + *Oxtr* Meadow: t = 2.4798, p = 0.0296; *Drd1* + *Drd2* + *Oxtr* Prairie vs *Drd1* + *Drd2* + *Oxtr* Meadow: t = 2.3769, p = 0.0310). Meadow voles have greater levels of *Drd2* only cells and *Drd1* + *Drd2* cells as compared to prairie voles across all subregions examined (Fig 2A: *Drd2* only Prairie vs *Drd2* only Meadow: t = −7.3465, p < 0.0001; *Drd1* + *Drd2* Prairie vs *Drd1* + *Drd2* Meadow: t = −6.9505, p < 0.0001; Fig 2B: *Drd2* only Prairie vs *Drd2* only Meadow: t = −4.6578, p < 0.0001; *Drd1* + *Drd2* Prairie vs *Drd1* + *Drd2* Meadow: t = −5.2969, p < 0.0001; Fig 2C: *Drd2* only Prairie vs *Drd2* only Meadow: t = −9.3429, p < 0.0001; *Drd1* + *Drd2* Prairie vs *Drd1* + *Drd2* Meadow: t = −7.3221, p < 0.0001).

### 2.3. *Oxtr* expression in dopamine receptor cell classes is largely unaffected by sociosexual experience

We next investigated whether mating influenced receptor distribution in either species. Partner preference testing confirmed species-specific differences social behaviors. Prairie voles spent more time huddling with their partner than with a novel conspecific (Fig 3B: 2-way ANOVA: main effect of conspecific: F(_1,13_) = 10.24, p = 0.007; main effect of sex: F(_1, 13_) = 0.3043, p = 0.8560; interaction [conspecific x sex]: F(_1,13_) = 0.2546, p = 0.6223), while meadow voles did not display a partner preference (Fig 3C). Overall differences in sociality between species were also evident in the amount of time spent in social chambers, with the unexpected finding that male meadow voles spent significantly more time in a social chamber compared to female meadow voles (Fig 3B: unpaired t-test: t = 3.034, p = 0.0126). Animals indicated by solid gray triangles or circles are those used in *in situ* experiments (Fig 3B and 3C).

**Figure 3.**
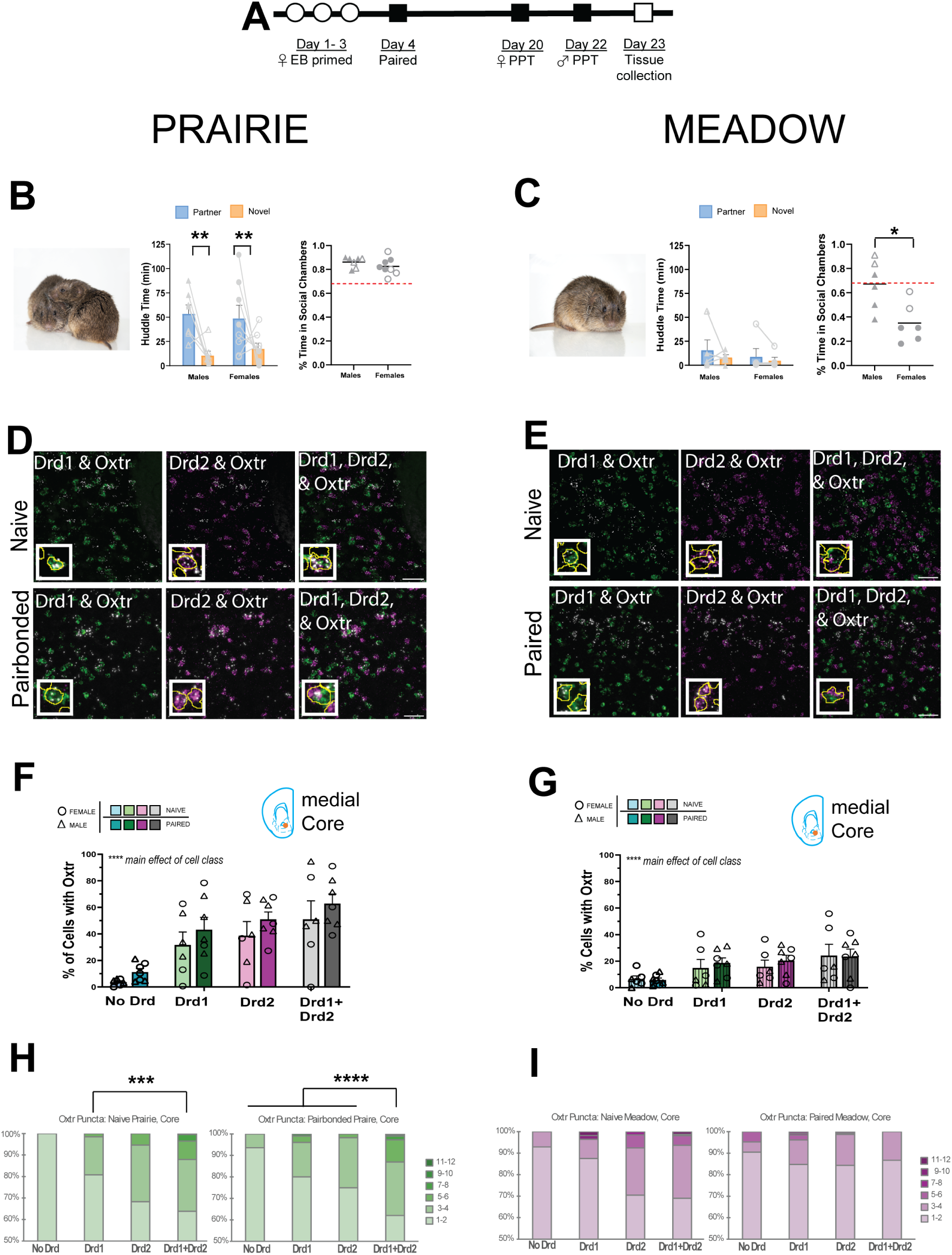
*Oxtr* distribution in dopamine receptor cell classes does not vary based on sociosexual experience. **A.** Experimental timeline for pairing and behavioral test for paired prairie and meadow vole cohorts. **B-C.** A partner preference test confirmed pair bond formation in prairie voles, but not meadow voles. Male and female prairie voles, and male meadow voles more time spent in social chambers compared to female meadow voles. Solid gray triangles and circles indicate animals used in *in situ* experiments. **D-E.** Representative images demonstrating co-labeling of *Oxtr* with *Drd1*, or *Drd2*, or both *Drd1* and *Drd2* in the nucleus accumbens core occurs in all cohorts: sexually naïve prairie voles (**D**: top row), pairbonded prairie voles (**D**: bottom row), sexually naïve meadow voles (**E**: top row) and paired meadow voles (**E**: bottom row). Image inset represents 40x image with 4x zoom. Scale bar = 50 µm. **F-G**. Prairie and meadow voles showed a main effect of cell class but no main effect of bond status on the percentage of cells with *Oxtr* across dopamine receptor cell class, indicating that this metric is not influenced by sociosexual experience but does scale as a function of cell dopamine receptor identity in both species. **H-I**. The number of *Oxtr* puncta was counted, and the frequency of puncta number was calculated in each dopamine receptor cell class. Naïve prairie voles had a greater puncta density in *Drd1+Drd2* cells compared to *Drd1* cells (**H**: left). Pairbonded prairie voles had a greater puncta density in *Drd1+Drd2* cells compared to all other dopamine receptor cell classes (**H**: right). Naïve meadow voles (**I**: left) and paired meadow voles (**I**: right) had no statistically significant differences in *Oxtr* puncta frequency across dopamine receptor cell classes. Error bars show SEM. n = 4-8 per group. * p < 0.05, ** p < 0.01, *** p < 0.001, **** p < 0.0001.

Mating had no effect on *Oxtr* distribution across dopamine receptor cell classes in either species, a pattern consistent across all subregions examined (Fig 3F-G; Supplementary Fig. 1A-B, E-F). Classification of the same 8 transcript combinations as in Figure 2 revealed that cell ratios do not differ in prairie voles before and after bonding, except for two differences: naïve prairie voles had greater no transcript expression in the core (Supplementary Fig 2A: t = 3.4716, p = 0.0061) and greater *Drd1+Drd2* expression in the medial shell (Supplementary Fig 2B: t = −4.0282, p = 0.0009) (all non-significant results in Table S1). Somewhat surprisingly, meadow voles showed more pronounced mating-induced differences, especially in the lateral shell. Naïve meadow voles exhibited greater *Drd2* only expression in the core and lateral shell (Supplementary Fig 2A: t = - 3.8622, p = 0.0016; Fig 2C: t = −3.0634, p = 0.0056, respectively), while *Drd1+Drd2* expression was higher in naïve animals in the medial and lateral shell (t = −3.6401, p = 0.0035; t = −3.0945, p = 0.0056). Interestingly, naïve meadow voles had greater *Drd2+Oxtr* and *Drd1+Drd2+Oxtr* cells compared to paired counterparts, though absolute levels in both groups remained low (<3% and <4%, respectively; Supplementary Fig 2C: t = −2.8353, p = 0.0089; t = −5.0947, p < 0.0001). Additionally, paired meadow voles had significantly more *Drd1*-only cells than naïve meadow voles (Fig 2C: t = 3.0992, p = 0.0056).

To gain a more nuanced assessment of relative transcript abundance in different cell classes, we counted fluorescent puncta per cell, which was feasible for *Oxtr* due to its low abundance. Across sexually naïve and mated prairie voles, more *Oxtr* puncta were observed in *Drd1*+*Drd2+* positive cells. (Fig 3H: Naïve-Generalized Fisher’s Exact Test: p = 0.0003; Post hoc Fisher’s Exact Test with Bonferroni correction: *Drd1* vs *Drd1*+*Drd2*: p = 0.0009; Pairbonded - Chi Squared Test of Independence: χ2 = 28.9829, p < 0.0001; Post hoc Chi Squared Test for Trend with Bonferroni correction: No Drd vs *Drd1*+*Drd2*: χ2 = 20.1709, p < 0.0001; *Drd1* vs *Drd1*+*Drd2*: χ2 = 30.3492, p < 0.0001; *Drd2* vs *Drd1*+*Drd2*: χ2 = 18.2546, p < 0.0001). Similar trends were observed in the medial and lateral shell of naïve and pairbonded prairie voles (Supplementary Fig. 1C & 1G; see detailed statistics table).

In contrast, meadow voles had lower *Oxtr* puncta counts, with no significant differences in puncta frequency across *Drd* receptor cell classes in either naïve or paired conditions (Fig 3I: naïve meadow: Generalized Fisher’s Exact Test: p = 0.0951; paired meadow: Generalized Fisher’s Exact Test: p = 0.1938). Similar patterns were observed in the medial and lateral shell of meadow voles (Supplementary Fig. 1D & 1 H; see Table S1).

### 2.4. Mapping of oxytocin and dopamine receptor expression across cell types in the prairie vole NAc

Finally, we used existing RNA sequencing datasets from the prairie vole NAc to comprehensively map *Oxtr* and *Drd* transcripts in the prairie vole NAc[28]. Tissue level sequencing revealed that *Drd1* and *Drd2* are expressed 10-20x higher levels than *Drd3*, *Drd4*, *Drd5*, and *Oxtr*. This pattern aligns with our FISH findings, which also indicate a higher relative abundance of *Drd1* and *Drd2* compared to *Oxtr*. (Fig. 4A). In addition, we observed that *Drd3* and *Drd5* transcripts are expressed in the NAc at levels commensurate with that of *Oxtr*, a gene with a well-established role in bonding.

**Figure 4.**
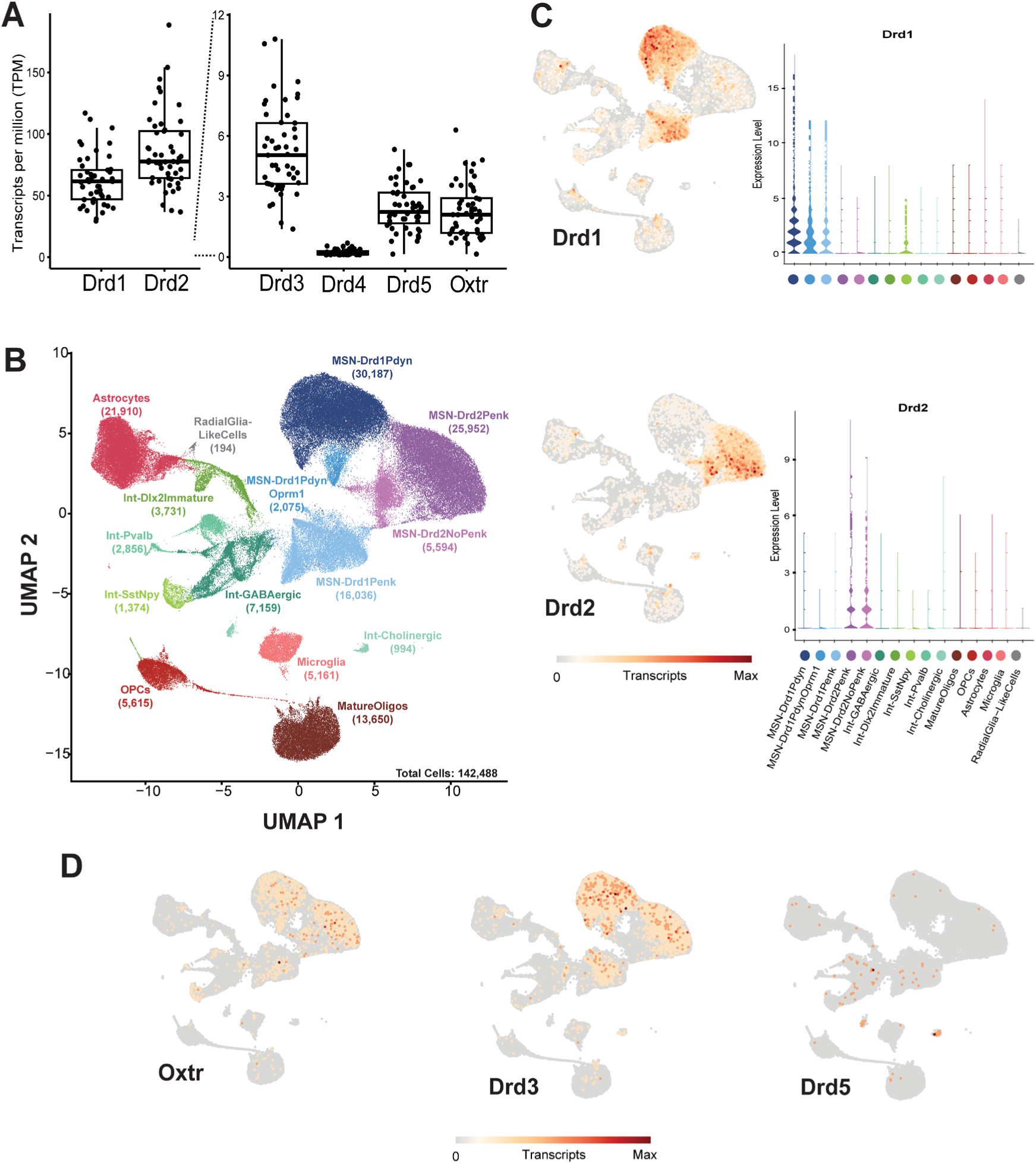
Mapping of oxytocin and dopamine receptor expression across cell types in the prairie vole nucleus accumbens. **A.** *Oxtr* and *Drd* transcript expression levels from prairie vole nucleus accumbens. *Drd1* and *Drd2* transcripts are expressed at high levels, whereas *Drd3, Drd4, Drd5,* and *Oxtr* transcript levels are significantly less abundant. **B.** Single-nucleus RNA sequencing data from prairie voles identifying 15 molecularly-defined cell types in the nucleus accumbens. **C-D**. Visualization of *Oxtr* and *Drd* transcript expression across all nucleus accumbens cell types (UMAP and violin plots). Single-nucleus RNA sequencing results support our FISH results in that *Oxtr* is expressed in both *Drd1* and *Drd2* cells, as well as in some astrocytes, mature oligodendrocytes, and all interneurons populations identified (somatostatin/neuropeptide Y, parvalbumin and GABAergic interneurons). *Drd1* and *Drd2* transcripts are expressed in medium spiny neuron populations (MSN’s) as well as astrocytes, oligodendrocytes, microglia, and various interneuron populations. *Drd5* transcripts appear selectively enriched in GABAergic and cholinergic interneurons, whereas *Drd3* transcripts are expressed across several cell types including MSN’s and sparsely across any other cell types.

To further investigate receptor expression in molecularly defined cell types in the NAc, we examined transcript expression in a single-nucleus RNA sequencing dataset from 39 prairie voles. This dataset was used to identify 15 primary cell types in the NAc, characterized by the expression of key marker genes, including *Drd1* and *Drd2* (Fig 4B)[27]. However, low-abundance transcripts such as *Oxtr*, *Drd3*, *Drd5*, suffer from high-levels of dropout (e.g. a failure to detect the transcript or false negative)[29,30]. This was evident when comparing *Oxtr* detection rates:1% via snRNA-seq vs 34-38% (depending on the NAc subregion) via FISH. Thus, while mapping low-abundance transcripts onto UMAP projections provides insights into which cell types express a given transcript, it does not accurately estimate the proportion of expressing cells.

To better visualize *Oxtr* and *Drd* transcript distribution across cell types, we plotted transcript-positive cells on a UMAP. This approach overlays transcript-expressing cells on top of the entire cell cluster. To provide a more accurate representation of expression levels for each cell type, we also plotted transcript counts per cell cluster (Fig 4C). Using this approach, we found that *Drd1* and *Drd2* were enriched in subsets of medium spiny neurons (MSN’s), but also detectable across all identified cell types (Fig. 4C). Due to high dropout rates associated with their low abundance, quantitative estimates of *Drd3*, *Drd5*, and *Oxtr* expression remain inconclusive. Despite dropout effects, UMAP projections suggest that *Oxtr* is expressed in MSNs and Sst-Npy interneurons, *Drd3* is localized to various MSN subtypes, and *Drd5* is potentially expressed in GABAergic and cholinergic interneurons (Fig. 4D). These distributions suggest potential cell-type specific roles for *Oxtr*, *Drd3* and *Drd5* signaling in modulating cell functions and behavior.

## 3. Discussion

This study presents the first detailed comparison of accumbal *Oxtr*, *Drd1*, and *Drd2* expression at the cellular level in sexually naïve and mated prairie and meadow voles. We demonstrate that the previously reported difference in *Oxtr* and OXTR expression between these two species (prairie > meadow)[6–9] is due, at least in part, to differences in the total number of cells expressing *Oxtr* transcripts in the NAc. Further, *Oxtr* expression occurs across *Drd* cell classes in both species, and this distribution does not differ by sex or as a function of mating/bonding. *Oxtr* puncta are enriched in *Drd1+Drd2* co-expressing cells, especially in prairie voles, suggesting preferential recruitment of these cells by oxytocin. Finally, we provide the first expression atlas for *Oxtr* and all known *Drd* receptors across molecularly defined cell types in the prairie vole NAc.

We note that while prairie voles exhibit a higher prevalence of OXTR expression overall, the distribution of *Oxtr* across *Drd*-expressing cell classes appears to be similar across species. This suggests that species differences in OXTR signaling occurs via a greater proportion of cells overall rather than preferentially engaging a specific DRD cell class in prairie voles. Given that oxytocin is released upon mating in prairie voles [31], this widespread activation may function as a filtering mechanism that enhances the saliency of mating-related sensory and reward signals, thereby reinforcing both the mating experience and the identity of the partner. Similar mechanisms have been proposed in other brains regions where OXTR-mediated modulation of local signal-to-noise ratios can selectively enhance relevant stimuli [32]. While this remains a working model, it provides a plausible cellular-level framework for how species differences in OXTR prevalence may contribute to the formation of enduring social bonds.

Notably, we observed that prairie voles had a higher number of *Oxtr*-expressing and co-expressing cells, particularly in the lateral shell of the NAc. This is especially intriguing, as most research has focused on the medial shell and core, leaving the lateral shell largely unexplored in the context of social behavior and pair bonding. Our findings highlight significant species differences in *Oxtr* expression and its co-localization with *Drd* receptors in the lateral shell, contributing to the growing literature on NAc heterogeneity [33–35]. These results suggest that the lateral shell may play a previously underappreciated role in social bonding, particularly in monogamous species by integrating oxytocin and dopamine signaling in a unique way.

While prior studies have reported increased *Drd1* and DRD1 levels following pair bonding in prairie voles[18], these studies have not provided cellular-level resolution. Our findings suggest that increased levels may result from upregulation of *Drd1* within existing populations rather than recruitment of new *Drd1+* cells. This distinction is important, as it implies that dopaminergic plasticity in bonding likely occurs through receptor upregulation in cells already expressing the receptor rather than new expression in cells without a history of *Drd1* expression. This suggests an enhancement of Drd1 signaling in a specific circuit rather than engagement of new circuits mediating bonding-dependent effects on behavior, although this needs to be experimentally tested in future work.

Our results provide particular relevance for understanding potential oxytocin:dopamine interactions at a cellular level. *Oxtr* expression within *Drd2+* cell classes creates an opportunity for heterodimer formation. *In vitro* studies using bioluminescence resonance energy transfer (BRET) demonstrated formation of D2R-Oxtr heterodimers, suggesting a direct molecular interaction between these receptors[36]. In these heterodimer-specific experiments, oxytocin activation enhanced D2R signaling and increased inhibition of the cAMP-PKA-CREB pathway[37]. Separately, radioligand binding experiments using ^3^H-raclopride showed that oxytocin increased D2R binding, suggesting an additional mechanism by which oxytocin modulates D2R function [38]. Together, these findings suggest that oxytocin may enhance D2 receptor function via two mechanisms: (1) direct heteromeric interactions that increase D2R affinity for dopamine, thereby amplifying inhibitory Gi/o signaling and reducing cAMP levels, and (2) increased receptor availability, potentially via upregulation of receptor density or altered receptor conformation. Both mechanisms would converge on enhanced D2R-mediated inhibition of the cAMP-PKA-CREB pathway, ultimately suppressing CREB-dependent transcription of genes involved in synaptic plasticity and reward processing such as c-Fos, TH, BDNF, and ΔFosB, all of which are known to contribute to pair bonding [39–44]. Of course, an important question remains: do these receptors merely colocalize within the same cells or do they form true heterodimers *in vivo*? Future research should explore the dynamic interactions between these receptor systems using super-resolution imaging or functional assays to address this question.

Additionally, our data revealed that *Oxtr* is expressed on a subset of cells positive for both *Drd1* and *Drd2,* and we observed increased *Oxtr* puncta density in these *Drd1+Drd2* cells, suggesting that these cells may exhibit enhanced oxytocinergic sensitivity. Overall, our data are consistent with prior work showing that 10-30% of NAc MSNs in other species exhibit dopamine receptor co-expression[45–48]. Notably, coordinated activation of multiple receptors (*Oxtr, Drd1, Drd2*) in the same cell may engage distinct intracellular signaling pathways that regulate bond-relevant transcriptional programs and/or neuronal activity. *Oxtr*, *Drd1*, and *Drd2* each activate distinct intracellular signaling cascades, yet they may also integrate signals to shape social behavior. OXTR is typically Gq-coupled, leading to phospholipase C (PLC) activation and intracellular calcium release, which modulates excitability and synaptic plasticity. In contrast, D2-like receptors (including DRD2) are Gi-coupled, inhibiting adenylate cyclase and reducing cAMP levels dampening excitability, while D1-like receptors (including DRD1) are Gs-coupled, stimulating adenylate cyclase and increasing cAMP production, promoting excitatory signaling [49–51]. Beyond their independent signaling roles, D1-D2 heteromers in rat NAc neurons and have been found to induce a unique pathway where the heteromer is Gq-coupled, stimulates the PLC pathway to release intracellular calcium from IP3 receptor-sensitive stores, leading to CaMKII activation and BDNF production. This mechanism may underlie synaptic plasticity and reinforce socially motivated behaviors[46]. Given the role of dopaminergic plasticity in social bonding, the co-expression of *Oxtr*, *Drd1*, and *Drd2* within the same cells--and the potential formation of heteromers—may enable complex intracellular cross-talk between these pathways, fine-tuning cellular responses to social stimuli. These interactions could shape neural circuits involved in pair bonding, and reinforce behaviors critical for maintaining social attachment[50,51].

The role of D1-D2 heteromers in reward processing extends beyond social bonding. Studies have demonstrated that repeated amphetamine (AMPH) administration leads to functional super sensitivity of the D1-D2 receptor heteromer, as evidenced by enhanced GTPγS binding, and also increases D1-D2 heteromer density in the NAc, as assessed by FRET techniques[52]. These findings are particularly relevant given that pair bonding in prairie voles protects against AMPH’s rewarding effects, with this buffering mediated via a D1-specific mechanism in males [53]. This suggests that pair bonding may modulate dopaminergic plasticity in a way that counteracts drug-induced neuroadaptations, highlighting a potential role for *Oxtr-Drd1-Drd2* co-expression in reshaping neural circuits involved in both natural and drug-related reward processing.

Finally, we provide a comprehensive map of *Drd* expression across molecularly-defined cell types in the prairie vole NAc, including for less-commonly studied dopamine receptors. Although *Drd3* and *Drd5* are expressed at lower levels in the NAc, evidence suggests they may play important roles in social behavior. DRD3 agonists disrupt social behavior and social recognition in rats[54,55], while dopamine DRD5 receptors modulate sexual behavior in mice[56], although these functions may be localized outside of the NAc. Furthermore, preliminary data, although underpowered, also suggests that DRD3, rather than DRD2 receptors, may be more important for bond formation in prairie voles[57]. Accordingly, our expression map offers a valuable reference for studying dopaminergic contributions to social behavior and underscores the need for receptor-specific investigations to clarify the role of DRD3 and DRD5 receptors in social bonding.

Finally, while we provide insight into accumbal *Oxtr*, *Drd1*, and *Drd2* expression and distribution at the cellular level in prairie and meadow voles, our study is not without its limitations. Our behavioral cohort sample sizes were relatively small, (6-9 animals/group). Also, because our sample includes prairie voles with different *Oxtr* genotypes, future studies should stratify animals by genotype to assess whether genetic variation contributes to species differences. Additionally, our workflow relied on a nuclear (DAPI) mask to identify transcript-positive cells, meaning transcripts outside the nucleus (e.g., in axon terminals or dendrites) were not captured. This is particularly relevant for Drd1, which has been shown to change with social experience at the mRNA level[58], but may not always translate to nuclear-proximal expression. Further, determining the relationship between the numbers of cells expressing a transcript and what this means for protein abundance and localization could provide further insight into their regulation[25]. More broadly, our findings highlight *Oxtr*-*Drd* co-expression as a potential mechanism for integrating oxytocin and dopamine signaling in the NAc, but the functional relevance of our observations has yet to be established.

In sum, our findings may have implications for understanding social deficits in neuropsychiatric conditions characterized by oxytocin-dopamine dysregulation, such as autism spectrum disorder (ASD) and schizophrenia. Given that disruptions in *Oxtr*-*Drd* interactions could impair social reward processing, future research should explore whether similar neuromodulatory mechanisms contribute to altered social motivation in these conditions. Investigating whether pharmacological modulation of *Oxtr* or *Drd* signaling in the NAc can rescue social deficits may provide a more targeted framework for therapeutic interventions. Beyond clinical applications, investigating the circuit-specific mechanisms through which oxytocin modulates social reward will be essential for bridging the gap between animal and human studies. A deeper understanding of these neuromodulatory interactions could refine our models of dopaminergic plasticity in social bonding and inform the development of target approaches for addressing social behavior impairments.

## 4. Methods

### 4.1. Animals

Prairie voles were bred in-house from colonies originating from the University of California Davis and Emory University, with all animals originally descended from wild animals collected in Illinois. Meadow voles were also bred in house from colonies originating from Smith College and University of California San Francisco. All voles were weaned at postnatal day 21. Animals were then housed in standard static rodent cages (17.5l x 9.0w. x 6.0h. in.) in groups of 2-4 with either same sex siblings or same sex voles from similar weaning time frames. Animals were given *ad libitum* access to water and rabbit chow (5326-3 by PMI Lab Diet). Rabbit chow was supplemented with sunflower seeds, dehydrated fruit bits, and alfalfa cubes. Cages were enriched with cotton nestlets, a plastic igloo, and a PVC pipe. Animals were kept in a temperature (23-26 °C) and humidity-controlled room with a 14:10 hour light-dark cycle.

All voles used in the studies ranged in ages of 62 - 104 days at the start of experiments and weighed between 28.4g – 48.4g. All procedures were approved under the University of Colorado’s Institute of Animal Care and Use Committee (IACUC) and performed in the light phase. All authors complied with the ARRIVE guidelines.

### 4.2. Sample Collection Timeline

All sexually naïve animals used in this study were housed in groups of 2-4 with same sex siblings or same sex voles from similar weaning time frames.

All paired animals used in this study consisted of one female and one male from different genetic backgrounds. Females were reproductively intact and were primed to induce estrus via subcutaneous injection of 20 ug/mL of estradiol benzoate (Cayman Chemical) every 24 hours for 3 days. On the third day of priming, females were paired with their designated partner and placed in a smaller standard rodent cage (11.0l x 6.5w. x 5.0h. in) with fresh bedding, food, water, and cage enrichment.

Sixteen days after pairing, we measured partner preference in female animals, and 18 days after pairing, we measured partner preference in male animals. Nineteen days after pairing, we extracted brains for analysis in 16 prairie vole pairs (8 pairs for a female focal animal and 8 pairs for a male focal animal) and 7 meadow vole pairs. Tissue from a subset of 4 animals per sex was used in *in situ* hybridization studies. These animals were selected based on partner preference score and total test huddle time. Specifically, all paired prairie voles used in *in situ* studies had a partner preference (> 66% time spent huddling with partner) and spent at >2% of total test time (216 seconds) huddling). The subset of prairie voles selected for the *in situ* studies were those with highest partner preference scores. All paired meadow voles used in *in situ* studies had no partner preference ((< 66% time spent huddling with partner) and/or <2% of total test time huddling).

### 4.3. Partner preference test

Partner preference tests (PPT) were performed as described in Scribner et al. 2020[59]. Briefly, partner and novel animals were tethered to opposite end walls of three-chamber plexiglass arenas (76.0 cm long, 20.0 cm wide, and 30.0 cm tall). Tethers consisted of an eye bolt attached to a chain of fishing swivels that slid into the arena wall. Animals were briefly anesthetized with isoflurane and attached to the tether using a zip tie around the animal’s neck. Two pellets of rabbit chow were given to each tethered animal and water bottles were secured to the wall within their access while tethered. After tethering the partner and novel animals, the experimental animal was placed in the center chamber of the arena. After a 10-minute acclimation period, opaque dividers between the chambers were removed, allowing the subject to move freely about the arena for three hours. Overhead cameras (Panasonic WVCP304) were used to video record tests.

The movement of all three animals in each test was scored using TopScan High-Throughput software v3.0 (Cleversys Inc) using the parameters from Ahern et al. [60] Behavior was analyzed using a Python script developed in-house (https://github.com/donaldsonlab/Cleversys_scripts) to calculate the following metrics: time spent in partner, novel, or center chamber, the time spent huddling with the partner/novel, and total locomotion.

### 4.4. Tissue Extraction & Brain Sectioning

Animals were sedated with isoflurane and euthanized via rapid decapitation. Brains were immediately extracted, rinsed in sterile saline, and then frozen on powdered dry ice. Samples were stored at −80° C for a minimum of 24 hours. 90 – 120 minutes before cryostat slicing, brains were moved to a −20° C freezer to acclimate to the temperature of the cryostat. Brain tissue was sliced coronally at 12 µm and nucleus accumbens tissue was collected between bregma 1.54mm to 0.74mm as per *The Mouse Brain Atlas in Stereotaxic Coordinates*[61] and mounted on Superfrost Plus slides (Fisherbrand). During slicing, slides were kept inside the cryostat to maintain a stable temperature and minimize RNA degradation. After slicing, tissues slices were stored at −20° C for 2 hours to support better adhesion of tissue to the slide and then moved to −80° C for subsequent storage.

### 4.5. Assessment of pregnancy status

After rapid decapitation, females were examined for embryos by dissection. The uterus was identified, and number of embryos was based on the number of independent protrusions.

### 4.6. Fluorescent RNAscope in situ Hybridization

Fluorescent *in situ* hybridization triple labeling of oxytocin receptor (*Oxtr*), dopamine receptor 1 (*Drd1*), and dopamine receptor 2 (*Drd2*) mRNA expression in the nucleus accumbens was performed using RNAscope Multiplex Fluorescent v2 Kit (ACD Bio). Species specific (*Microtus orchrogaster*) probes were used for dopamine receptor 1 (Mo-*Drd1*, Catalog #: 588161-C3); dopamine receptor 2 (Mo-*Drd2*, Catalog #: 534471-C2); and oxytocin receptor (Mo-*Oxtr*, Catalog #: 500721); 3-plex positive control probes Mo-Pol2ra (polypeptide A 220kDa, Catalog # 563291), Ppib (peptidylprolyl isomerase B, Catalog # 533491), and Ubb (ubiquitin B, Catalog #300040); 3 plex negative control probe DapB (4-hydroxy-tetrahydrodipicolinate reductase from *Bacillus subtilis*, Catalog # 320871). Sections were fixed using 4% PFA in 1X PBS and then dehydrated in a series of ethanol washes. Endogenous peroxidases were blocked via *RNAscope* H_2_O_2_ and tissues were permeabilized using *RNAscope* protease III. Slides were hybridized with the probes in a HybEZ oven (ACD), at 40°C for 2 hours and then slides were subjected to signal amplification according to manufacturer’s instructions. Hybridization signals were detected using Perkin Elmer Opal Dyes 520 (Catalog #: fp1487001kt), 570 (Catalog #: fp1488001kt), and 650 (Catalog #: fp1496001kt) at a 1:50 dilution. Slides were counterstained with RNAscope DAPI for nuclear visualization and coverslipped with ProLong Diamond Antifade Mountant (Invitrogen, Catalog # P36970). Slides were stored covered at 4°C and imaged within 2 weeks.

### 4.7. Microscopy

Slices were imaged using a Nikon A1 Laser Scanning confocal Microscope at the University of Colorado Boulder MCDB Light Microscopy Core Facility (RRID:SCR_018993). Images were taken in one sitting of the anterior portion of the nucleus accumbens at Bregma 1.54mm as defined according to *The Mouse Brain Atlas in Stereotaxic Coordinates*[61] on both left and right hemispheres. Confocal images were taken with a 40x lens in four channels (blue, green, red, and infared) for each hemisphere of the core, and medial and lateral shell of the nucleus accumbens. Confocal stacks were projected as single images using maximum fluorescence and used for analysis.

### 4.8. Image Analysis

Images were analyzed using Fiji Image J software (version 2.14.0/1.54f) [62]. Images were split into 4 channels: channel 0 / DAPI, channel 1 / *Drd1*, channel 2 / *Drd2*, and channel 3 / *Oxtr*. Regions of interest (ROI’s) outlining DAPI nuclei labeling were automatically generated in Fiji Image J. Thresholds for DAPI nuclear staining were established by eye to eliminate background and provide an accurate overlay of nuclei in each image. Accuracy of DAPI masks was quantified for a subset of images (10%, 1 image per animal) comparing experimenter number of identified nuclei per image vs automatically identified number of nuclei per image. Number of nuclei identified by person vs machine did not differ by more than 5% and thus, automatically generated ROI’s for DAPI were used for all further analysis. This DAPI mask overlay was then applied to *Drd1*, *Drd2*, and *Oxtr* images (Fig. 1C). Images underwent a white top-hat transformation to enhance contrast and improve detection of localized bright features facilitating identification and quantification of puncta. Signal data for *Drd1*, *Drd2*, and *Oxtr* images was then collected including the region of interest number / nuclei number, minimum, mean and maximum intensity values, and % area for both 16- and 8-bit depth data. This final cellular distribution data was analyzed via custom-generated Python code (https://github.com/donaldsonlab/RNAscope paper) to identify the number of positively labeled nuclei for each channel, as well as co-expression in the form of double and triple labeled cells. *Oxtr* puncta were quantified using Morphological Filters Plugin (https://github.com/ijpb/MorphoLibJ) and Object Inspector (2D/3D) Plugin [63] in Fiji Image J.

### 4.9. Methods for bulk RNAsequencing

Animals were sacrificed by rapid decapitation and the NAc was dissected out of the brain prior to processing for RNA-Seq. Sequence mapping and counting was performed as described in Sadino et al. [28] (GEO: GSE192661).

### 4.10. Methods for single nucleus RNAsequencing

Methods were as described in Brusman et al [27]. Briefly, single nuclei were isolated, and samples were sequenced using Chromium next GEM Single Cell 5’ kit v2 from 10X Genomics. Single nuclei RNA-seq (snRNA-seq) libraries were prepared according to the manufacturer’s instructions. Library quality was assessed using the Agilent High Sensitivity D5000 ScreenTape System and were subsequently sequenced using paired-end sequencing on an Illumina NovaSeq6000. Raw reads were aligned and counted using the Cellranger v3.1.0 analysis pipeline, and Seurat v4.3.0 in R v4.2.2 was used to analyze single nuclei data. GEO (GSE255620).

### 4.11. Statistical Analysis

Data are shown as means ± standard error of the mean. Statistical significance α was set at 0.05, unless otherwise specified. All n values represent number of animals. Depending on the statistical test, statistical analyses were carried out using GraphPad PRISM (version 10.4.1, Graphpad, San Diego, CA); R (v 4.2.3, (2023-03-15 ucrt)), or Python (v 3.12.4) in a reproducible computing environment. RStudio (v 2024.12.0.467, Boston, MA) was used as the integrated development environment (IDE) for R analyses, while Python scripts were executed in Juypter Notebook (v 7.0.8) via Anaconda Navigator (v 2.6.4). In R, the following packages were used: ggplot2 (v 3.4.2); ggpubr (v.0.6.0); TMB (v 1.9.11). In Python, the following packages were used: NumPy: 1.26.4; SciPy: 1.13.1;pandas: 2.2.2; matplotlib: 3.8.4; and seaborn: 0.13.2. Normality of the data was assessed using Shapiro-Wilk tests in GraphPad PRISM, and parametric tests were chosen accordingly. Outliers were tested for using the Robust Regression and Outlier Removal (ROUT) method in GraphPad Prism (10.4.1), with a False Discovery Rate (Q) set at 1%. Additional statistical tests including 2-way repeated measures ANOVA’s; post hoc Šidák’s multiple comparisons test; t-tests and error propagation and false discovery rate; generalized Fisher’s Exact Test with post hoc Fisher’s Exact Test with Bonferroni correction; Chi Square Test of Independence with post hoc Chi Squared Test for Trend with Bonferroni correction were all conducted to evaluate differences between groups. In cases where differences were not found between males and females, sexes were combined.

### 4.12. Oxtr SNP Genotyping

All prairie voles were genotyped post-mortem for nucleotide 213739 (NT213739), as this robustly predicts *Oxtr* expression in the nucleus accumbens of prairie voles, where CC>CT>TT[64]. DNA was extracted from toes using The Jackson Laboratory Quick DNA purification protocol. The *Oxtr* SNP genotyping assay was designed using IDT’s rhAmp SNP Genotyping technology (Design ID: CD.GT.GFDG0397.1, IDT) and performed on an Applied Biosystems QuantStudio 3 qPCR machines. Samples were genotyped in duplicate in MicroAmp Fast Optical 96 well reaction plate (Applied BioSystems) in a reaction containing 2 µL of DNA, 0.25 µL of probe rhAmp SNP Assay, 0.1 µL nuclease free H2O, 0.13 µL rhAmp Reporter Mix, and 2.52 µL rhAmp Genotyping Master Mix for a total volume of 5 µL per well. Genotyping was performed using the following primers: 5’GAATCATCCCACCGTGC; 5’ GGAATCATCCCACCGTGT; and GCGTCAGTCCCTTATCGACCT. The cycling conditions were those as determined by the manufacturer, specifically: 95°C for 10 min, 40 x (95°C for 10 s, 60°C for 30 s, and 68°C for 20s) and 99°C for 15 min.

## Data Availability

Data used in each figure have been deposited on GitHub and will be publicly available on Dryad as of the date of publication. Any additional information required to reanalyze the data reported in this paper is available from the lead contact upon request.

## Acknowledgements

We thank the voles for their sacrifice and contribution to research. We thank Jessica Abazaris and the rest of the animal care staff at the University of Colorado Boulder for their excellent care of the voles. Kelly Winther, Katie Gallagher, and Kresil Gordon managed the animal colony and provided experimental support. We acknowledge the Light Microscopy Core Facility, Porter B047, B049, B051 and B059 at the University of Colorado Boulder (RRID:SCR_018993) for help and advice with microscopy and thank Dr. James D. Orth for his assistance. We thank the Donaldson lab for their feedback and support. This work was supported by awards from the Dana Foundation, the Whitehall Foundation, National Science Foundation (NSF) IOS-1827790, and National Institute of Health (NIH) DP2OD026143 to Z.R.D. and T32 DA 17637 support to M.K.L.

## Author Contributions

Conceptualization: MKL, ZRD. Formal analysis: MKL, JCS, CAG, LEB, JMS, KEW, DSWP. Funding acquisition: MKL, ZRD. Investigation: MKL, JCS, CAG, LEB, JMS. Project management: ZRD. Resources: ZRD. Visualization: MKL, LEB, JMS; Writing – original draft: MKL, ZRD. Writing – review & editing: MKL, LEB, JMS, ZRD

## Additional Information Competing Interests

The authors declare no conflicts of interest or competing interests.

