## Supplementary Materials for "Oxytocin and Dopamine Receptor Expression: Cellular Level Implications for Pair Bonding"

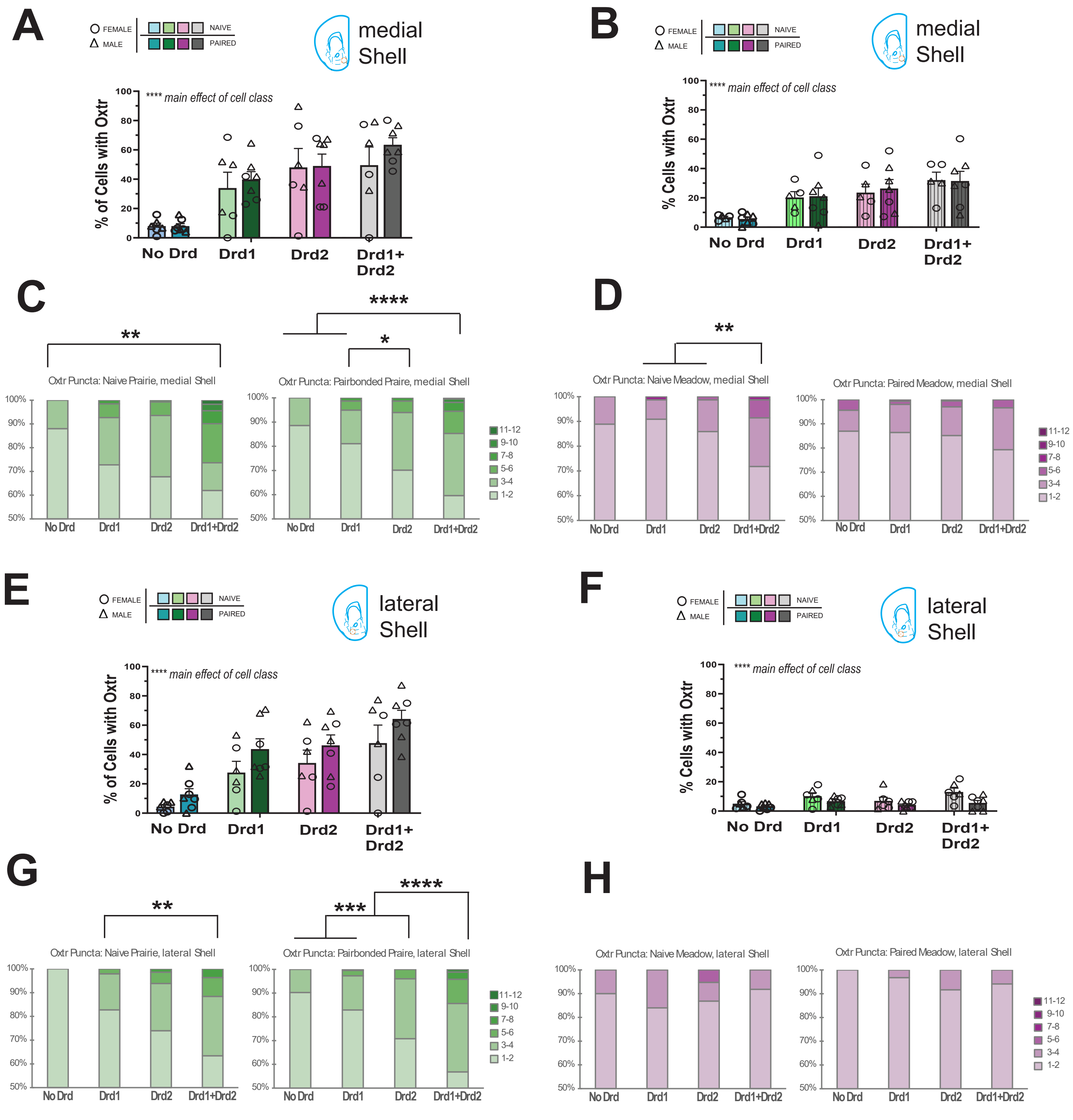


**Supplementary Figure 1. *Oxtr* distribution in dopamine receptor cell classes & *Oxtr* puncta frequency in the nucleus accumbens shell**

**A-B, E-F**. Percentage of cells with *Oxtr* across dopamine receptor cell classes in prairie voles (**A, E)** and meadow voles (**B, F**). In both the nucleus accumbens medial shell (**A-B**) and lateral shell (**E-F**), prairie and meadow voles showed a main effect of cell class, but no main effect of bond status on the percentage of cells with *Oxtr* across dopamine receptor cell class, indicating that this metric is not influenced by sociosexual experience.

**C-D**. In the medial shell, the number of *Oxtr* puncta was counted in each dopamine receptor cell class in sexually naïve prairie voles (**C**: right) and pairbonded prairie voles (**C**: left). Naïve prairie voles had a greater density of *Oxtr* puncta in *Drd1+Drd2* cells compared to No Drd cells. Pairbonded prairie voles had a greater density of *Oxtr* puncta in *Drd1+Drd2* cells compared to No Drd cells and *Drd1* cells. Additionally, *Drd2* cells had higher frequencies of *Oxtr* puncta compared to *Drd1* cells. In naïve meadow voles (**D**: right) there was a higher frequency of *Oxtr* puncta in *Drd1+Drd2* cells compared to *Drd1* or *Drd2* cells. Paired meadow voles (**D**: left) had no differences in *Oxtr* puncta frequency across the different cell classes.

**G-H**. In the lateral shell, naïve prairie voles had a greater density of *Oxtr* puncta in *Drd1+Drd2* cells compared to *Drd1* cells (**G**: left). Pairbonded prairie voles had a greater density of *Oxtr* puncta in *Drd1+Drd2* cells compared to all other cell classes. Additionally, *Drd2* cells had higher frequencies of *Oxtr* puncta compared to *Drd1* cells and No Drd cells (**G**: right)). Naïve meadow voles (**H**: left) and paired meadow voles (**H**: right) had no statistically significant differences in *Oxtr* puncta frequency across dopamine receptor cell classes.

Error bars show SEM. n = 6-8 per group. * $p < 0.05$, ** $p < 0.01$, *** $p < 0.001$, **** $p < 0.0001$.

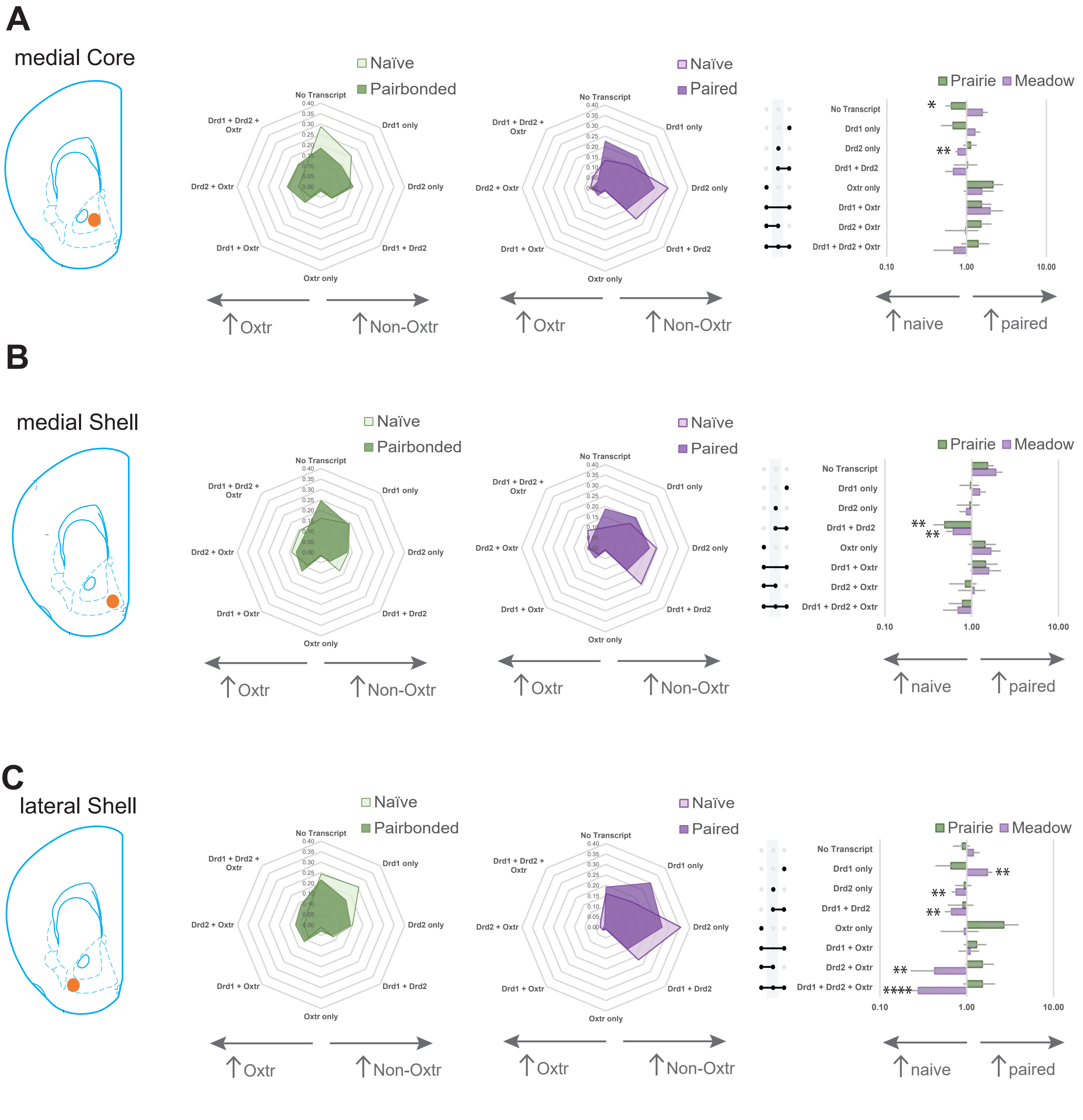


**Supplemental Figure 2. Sociosexual experience does not affect overall cell class receptor distribution in prairie voles.** **A-C.** Quantification of *Oxtr*, *Drd1*, and *Drd2* positive cells and their various co-expression combinations in the nucleus accumbens core (A), medial shell (B), and lateral shell (C) in prairie voles (Light green = sexually naïve. Dark green = pairbonded) and in meadow voles (Light purple = sexually naïve. Dark purple = paired). Fold change of paired to naive vole of each transcript category was quantified and compared between sociosexual experience for each subregion.

Pair bonded and sexually naïve prairie voles exhibit similar receptor expression across all cell classes and subregions, except for two differences: naïve prairie voles had greater no transcript expression in the core and greater *Drd1+Drd2* expression in the medial shell.

In meadow voles, sexually naïve and paired individuals showed more pronounced differences, particularly in the lateral shell. Naïve meadow voles exhibited greater *Drd2* only expression in both the core and lateral shell, while *Drd1+Drd2* expression was higher in naïve animals in the medial and lateral shell. Naïve meadow voles had fewer *Drd2+Oxtr* and *Drd1+Drd2+Oxtr* cells compared to paired counterparts, though absolute levels in both groups remained low (<3% and <4%, respectively). Additionally, paired meadow voles had significantly more *Drd1* only cells than naïve meadow voles.

Error bars show EP (error propagation). N = 6-8 per group. * $p < 0.05$, ** $p < 0.01$, *** $p < 0.001$, **** $p < 0.0001$.

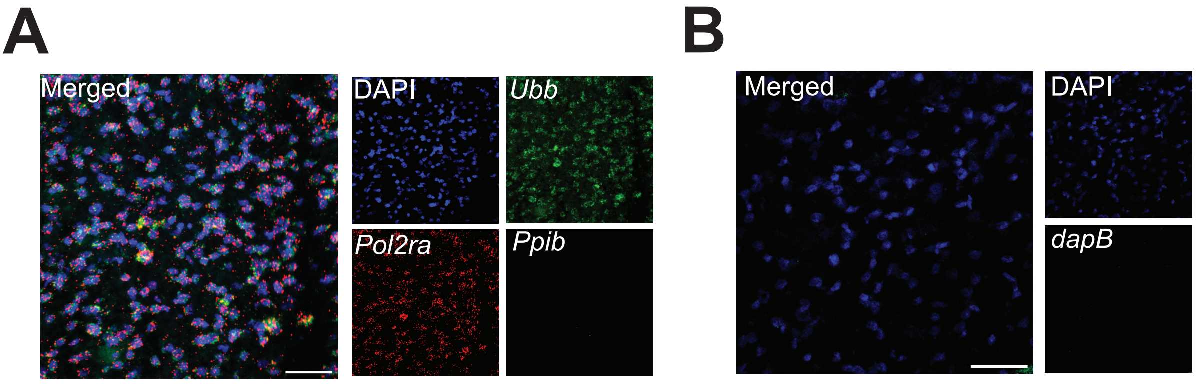


**Supplementary Figure 3. Experimental Controls**
**A**. Positive control probes to housekeeping genes *Ubb* (high abundance), *Pol2ra* (medium abundance), and *Ppib* (low abundance) show genes with varying degrees of expression and confirm assay success and probe visualization.
**B**. Negative control probe target to bacterial *dapB* confirms no nonspecific binding of fluorophores.
Images taken at 40x. Scale bars = 50 μm.

| Cohort | Animal # | Sex | Oxtr Genotype | Oxtr Phenotype |
|---|---|---|---|---|
| Sexually Naïve | 2721 | M | C/C | High |
| Sexually Naïve | 2773 | M | T/T | Low |
| Sexually Naïve | 2793 | M | C/T | Mid |
| Sexually Naïve | 2725 | F | C/T | Mid |
| Sexually Naïve | 2769 | F | C/T | Mid |
| Sexually Naive | 2790 | F | C/T | Mid |
| Pairbonded | 2763 | M | C/T | Mid |
| Pairbonded | 2770 | M | C/T | Mid |
| Pairbonded | 2776 | M | C/T | Mid |
| Pairbonded | 2782 | M | C/T | Mid |
| Pairbonded | 2794 | F | C/C | High |
| Pairbonded | 2812 | F | C/T | Mid |
| Pairbonded | 2816 | F | C/C | High |
| Pairbonded | 2829 | F | C/C | High |

**Supplementary Table 2. Oxtr SNP Genotypes in Prairie Voles**
