## Supplemental Table 1 for "Oxytocin and Dopamine Receptor Expression: Cellular Level Implications for Pair Bonding"

| Cohort Type | Region | Measurement | Statistical Test | Comparison | F/T/Chi-Sq Statistic | ° of freedom | p | * | 95% CI | Fig. | Stats Program Used |
|---|---|---|---|---|---|---|---|---|---|---|---|
| Prairie & Meadow Voles (all bond statuses) | Medial Core | % Positive Cells | 2 way RMANOVA | Interaction (Transcript Identity x Species) | 6.816 | 3.72 | 0.0004 | *** | | 1D | Prism |
| | | | | Species | 4.255 | 1, 24 | 0.0501 | ns | | | |
| | | | | Transcript Identity | 45.540 | 3.72 | <0.0001 | **** | | | |
| | | | | Subject | 0.843 | 24, 72 | 0.6717 | ns | | | |
| | | % Positive Cells | Post-hoc: Sidak's multiple comparisons | NT Prairie vs NT Meadow | | | 0.7768 | ns | | 1D | Prism |
| | | | | Drd1 Prairie vs Drd1 Meadow | | | >0.999 | ns | | | |
| | | | | Drd2 Prairie vs Drd2 Meadow | | | 0.3184 | ns | | | |
| | | | | Oxtr Prairie vs Oxtr Meadow | | | <0.0001 | **** | | | |
| Prairie & Meadow Voles (all bond statuses) | Medial Shell | % Positive Cells | 2 way RMANOVA | Interaction (Transcript Identity x Species) | 5.797 | 3, 69 | 0.0014 | ** | | 1E | Prism |
| | | | | Species | 0.972 | 1, 23 | 0.3345 | ns | | | |
| | | | | Transcript Identity | 54.390 | 3, 69 | <0.0001 | **** | | | |
| | | | | Subject | 0.904 | 23, 69 | 0.5935 | ns | | | |
| | | % Positive Cells | Post-hoc: Sidak's multiple comparisons | NT Prairie vs NT Meadow | | | 0.7453 | ns | | 1E | Prism |
| | | | | Drd1 Prairie vs Drd1 Meadow | | | >0.9999 | ns | | | |
| | | | | Drd2 Prairie vs Drd2 Meadow | | | 0.0614 | ns | | | |
| | | | | Oxtr Prairie vs Oxtr Meadow | | | 0.0041 | ** | | | |
| Prairie & Meadow Voles (all bond statuses) | Lateral Shell | % Positive Cells | 2 way RMANOVA | Interaction (Transcript Identity x Species) | 11.490 | 3,72 | <0.0001 | **** | | 1F | Prism |
| | | | | Species | 10.670 | 1, 24 | 0.0033 | ** | | | |
| | | | | Transcript Identity | 49.880 | 3, 72 | <0.0001 | **** | | | |
| | | | | Subject | 0.716 | 24, 72 | 0.8194 | ns | | | |
| | | % Positive Cells | Post-hoc: Sidak's multiple comparisons | NT Prairie vs NT Meadow | | | 0.9463 | ns | | 1F | Prism |
| | | | | Drd1 Prairie vs Drd1 Meadow | | | 0.8534 | ns | | | |
| | | | | Drd2 Prairie vs Drd2 Meadow | | | 0.1460 | ns | | | |
| | | | | Oxtr Prairie vs Oxtr Meadow | | | <0.0001 | **** | | | |
| Prairie & Meadow Voles (all bond statuses) | Medial Core | % Positive Cells, all 8 transcript combinations | t-test on a ratio of means with error propagation & FDR corrected p-value | NT Prairie vs NT Meadow | 1.3679 | | 0.2318 | | 0.8784, 1.5995 | 2A | Python |
| | | | | Drd1 only Prairie vs Drd1 only Meadow | -0.7736 | | 0.4410 | | 0.5801, 1.1843 | | |
| | | | | Drd2 only Prairie vs Drd2 only Meadow | -7.3465 | | 0.0000 | **** | 0.4266, 0.6705 | | |
| | | | | Oxtr only Prairie vs Oxtr only Meadow | 1.0077 | | 0.3612 | | 0.6325, 2.1261 | | |
| | | | | D1/D2 Prairie vs D1/D2 Meadow | -6.9505 | | 0.0000 | **** | 0.2649, 0.5913 | | |
| | | | | D1/Oxtr Prairie vs D1/Oxtr Meadow | 2.1637 | | 0.0774 | | 1.1121, 3.5861 | | |
| | | | | D2/Oxtr Prairie vs D2/Oxtr Meadow | 2.095 | | 0.0774 | | 1.0626, 3.3039 | | |
| | | | | D1/D2/Oxtr Prairie vs D1/D2/Oxtr Meadow | 1.9138 | | 0.0936 | | 0.9531, 3.6020 | | |
| Prairie & Meadow Voles (all bond statuses) | Medial Shell | % Positive Cells, all 8 transcript combinations | t-test on a ratio of means with error propagation & FDR corrected p-value | NT Prairie vs NT Meadow | 1.6107 | | 0.1472 | | 0.9052, 1.9133 | 2B | Python |
| | | | | Drd1 only Prairie vs Drd1 only Meadow | 0.026 | | 0.9793 | | 0.7209, 1.2865 | | |
| | | | | Drd2 only Prairie vs Drd2 only Meadow | -4.6578 | | 0.0000 | **** | 0.3821, 0.7512 | | |
| | | | | Oxtr only Prairie vs Oxtr only Meadow | 1.6822 | | 0.1472 | | 0.8852, 2.3948 | | |
| | | | | D1/D2 Prairie vs D1/D2 Meadow | -5.2969 | | 0.0000 | **** | 0.2850, 0.6747 | | |
| | | | | D1/Oxtr Prairie vs D1/Oxtr Meadow | 1.8161 | | 0.1472 | | 0.9369, 2.9779 | | |
| | | | | D2/Oxtr Prairie vs D2/Oxtr Meadow | 1.6505 | | 0.1472 | | 0.8754, 2.2372 | | |
| | | | | D1/D2/Oxtr Prairie vs D1/D2/Oxtr Meadow | 0.9576 | | 0.3892 | | 0.7022, 1.8533 | | |
| Prairie & Meadow Voles (all bond statuses) | Lateral Shell | % Positive Cells, all 8 transcript combinations | t-test on a ratio of means with error propagation & FDR corrected p-value | NT Prairie vs NT Meadow | 1.5862 | | 0.1159 | | 0.9281, 1.6449 | 2C | Python |
| | | | | Drd1 only Prairie vs Drd1 only Meadow | -0.826 | | 0.4106 | | 0.5377, 1.1904 | | |
| | | | | Drd2 only Prairie vs Drd2 only Meadow | -9.3429 | | 0.0000 | **** | 0.3581, 0.5830 | | |
| | | | | Oxtr only Prairie vs Oxtr only Meadow | 1.6959 | | 0.0930 | | 0.7351, 4.3891 | | |
| | | | | D1/D2 Prairie vs D1/D2 Meadow | -7.3221 | | 0.0000 | **** | 0.2698, 0.5811 | | |
| | | | | D1/Oxtr Prairie vs D1/Oxtr Meadow | 4.4565 | | 0.0000 | **** | 3.3782, 7.1947 | | |
| | | | | D2/Oxtr Prairie vs D2/Oxtr Meadow | 2.4798 | | 0.0148 | * | 1.9154, 9.2412 | | |
| | | | | D1/D2/Oxtr Prairie vs D1/D2/Oxtr Meadow | 2.3769 | | 0.0194 | * | 1.6882, 8.6376 | | |
| Prairie Voles (male & female) | n/a | Huddle time | 2 way ANOVA | Conspecific x Sex | 0.2546 | 1, 13 | 0.6223 | ns | | 3B | Prism |
| | | | | Conspecific | 10.2400 | 1, 13 | 0.0070 | ** | | | |
| | | | | Sex | 0.0343 | 1, 13 | 0.8560 | ns | | | |
| | | | | Subject | 0.2677 | 13, 13 | 0.9879 | ns | | | |
| | | | | Conspecific x Sex | 0.7936 | 1, 8 | 0.7936 | ns | | | |

| | | | | | | | | | | | |
|---|---|---|---|---|---|---|---|---|---|---|---|
| Meadow Voles (male & female) | n/a | Huddle time | 2 way ANOVA | Conspecific | 0.9002 | 1, 8 | 0.3705 | ns | | 3C | Prism |
| | | | | Sex | 0.4010 | 1, 8 | 0.5442 | ns | | | |
| | | | | Subject | 1.4060 | 6, 6 | 0.3448 | ns | | | |
| Prairie Voles | n/a | % time in social chamber | unpaired t-test | Male Prairie vs Female Prairie | 1.172 | 13 | 0.26 | ns | | 3B | Prism |
| Meadow Voles | n/a | % time in social chamber | unpaired t-test | Male Meadow vs Female Meadow | 3.0340 | 10 | 0.0126 | * | | 3C | Prism |
| Prairie: Naïve & Pairbonded | Medial Core | % Cells with Oxtr | 2 way RMANOVA | Interaction (Cell Class x Bond status) | 0.112 | 3,33 | 0.9524 | ns | | 3F | Prism |
| | | | | Bond status | 1.103 | 1, 11 | 0.3162 | ns | | | |
| | | | | Cell Class | 41.190 | 3, 33 | <0.0001 | **** | | | |
| | | | | Subject | 9.904 | 11, 33 | <0.0001 | **** | | | |
| | | % Cells with Oxtr | Post-hoc: Sidak's multiple comparisons | NT Naive vs NT Pairbonded | | | 0.9485 | ns | | 3F | Prism |
| | | | | Drd1 Naïve vs Drd1 Pairbonded | | | 0.8002 | ns | | | |
| | | | | Drd2 Naïve vs Drd2 Pairbonded | | | 0.7631 | ns | | | |
| | | | | Oxtr Naïve vs Oxtr Pairbonded | | | 0.7830 | ns | | | |
| Prairie: Naïve & Pairbonded | Medial Shell | % Cells with Oxtr | 2 way RMANOVA | Interaction (Cell Class x Bond status) | 0.925 | 3, 33 | 0.4397 | ns | | SF 1A | Prism |
| | | | | Bond status | 0.278 | 1, 11 | 0.6117 | ns | | | |
| | | | | Cell Class | 41.130 | 3, 33 | <0.0001 | **** | | | |
| | | | | Subject | 8.937 | 11, 33 | <0.0001 | **** | | | |
| | | % Cells with Oxtr | Post-hoc: Sidak's multiple comparisons | NT Naive vs NT Pairbonded | | | >0.9999 | ns | | SF 1A | Prism |
| | | | | Drd1 Naïve vs Drd1 Pairbonded | | | 0.9774 | ns | | | |
| | | | | Drd2 Naïve vs Drd2 Pairbonded | | | >0.9999 | ns | | | |
| | | | | Oxtr Naïve vs Oxtr Pairbonded | | | 0.6492 | ns | | | |
| Prairie: Naïve & Pairbonded | Lateral Shell | % Cells with Oxtr | 2 way RMANOVA | Interaction (Cell Class x Bond status) | 0.304 | 3, 33 | 0.8222 | ns | | SF 1E | Prism |
| | | | | Bond status | 2.408 | 1, 11 | 0.1490 | ns | | | |
| | | | | Cell Class | 32.550 | 3, 33 | <0.0001 | **** | | | |
| | | | | Subject | 6.038 | 11, 33 | <0.0001 | **** | | | |
| | | % Cells with Oxtr | Post-hoc: Sidak's multiple comparisons | NT Naive vs NT Pairbonded | | | 0.8926 | ns | | SF 1E | Prism |
| | | | | Drd1 Naïve vs Drd1 Pairbonded | | | 0.4270 | ns | | | |
| | | | | Drd2 Naïve vs Drd2 Pairbonded | | | 0.6932 | ns | | | |
| | | | | Oxtr Naïve vs Oxtr Pairbonded | | | 0.4050 | ns | | | |
| Meadow: Naïve & Paired | Medial Core | % Cells with Oxtr | 2 way RMANOVA | Interaction (Cell Class x Bond status) | 0.564 | 3, 33 | 0.6427 | ns | | 3G | Prism |
| | | | | Bond status | 0.079 | 1, 11 | 0.7838 | ns | | | |
| | | | | Cell Class | 14.320 | 3, 33 | <0.0001 | **** | | | |
| | | | | Subject | 10.420 | 11, 33 | <0.0001 | **** | | | |
| | | % Cells with Oxtr | Post-hoc: Sidak's multiple comparisons | NT Naive vs NT Paired | | | >0.9999 | ns | | 3G | Prism |
| | | | | Drd1 Naïve vs Drd1 Paired | | | 0.9728 | ns | | | |
| | | | | Drd2 Naïve vs Drd2 Paired | | | 0.9428 | ns | | | |
| | | | | Oxtr Naïve vs Oxtr Paired | | | >0.999 | ns | | | |
| Meadow: Naïve & Paired | Medial Shell | % Cells with Oxtr | 2 way RMANOVA | Interaction (Cell Class x Bond status) | 0.1684 | 3, 30 | 0.9168 | ns | | SF 1B | Prism |
| | | | | Bond status | 0.004669 | 1, 10 | 0.9469 | ns | | | |
| | | | | Cell Class | 25.25 | 3, 30 | <0.0001 | **** | | | |
| | | | | Subject | 9.224 | 10, 30 | <0.0001 | **** | | | |
| | | % Cells with Oxtr | Post-hoc: Sidak's multiple comparisons | NT Naive vs NT Paired | | | 0.9998 | ns | | SF 1B | Prism |
| | | | | Drd1 Naïve vs Drd1 Paired | | | >0.9999 | ns | | | |
| | | | | Drd2 Naïve vs Drd2 Paired | | | 0.9935 | ns | | | |
| | | | | Oxtr Naïve vs Oxtr Paired | | | >0.9999 | ns | | | |
| Meadow: Naïve & Paired | Lateral Shell | % Cells with Oxtr | 2 way RMANOVA | Interaction (Cell Class x Bond status) | 1.686 | 3, 33 | 0.1889 | ns | | SF 1F | Prism |
| | | | | Bond status | 4.548 | 1, 11 | 0.0563 | ns | | | |
| | | | | Cell Class | 6.242 | 3, 33 | 0.0018 | ** | | | |
| | | | | Subject | 3.529 | 11, 33 | 0.0024 | ** | | | |
| | | % Cells with Oxtr | Post-hoc: Sidak's multiple comparisons | NT Naive vs NT Paired | | | 0.9180 | ns | | SF 1F | Prism |
| | | | | Drd1 Naïve vs Drd1 Paired | | | 0.5287 | ns | | | |
| | | | | Drd2 Naïve vs Drd2 Paired | | | 0.7238 | ns | | | |
| | | | | Oxtr Naïve vs Oxtr Paired | | | 0.0158 | * | | | |
| | | | Generalized Fisher's Exact Test | No Drd vs Drd 1 vs Drd2 vs Drd1+Drd2 | na | | 0.0003 | *** | | | |
| | | | | No Drd vs Drd1 | na | | 1.0000 | ns | | | |
| | | | | No Drd vs Drd2 | na | | 0.2674 | ns | | | |

| | | | | | | | | | | | |
|---|---|---|---|---|---|---|---|---|---|---|---|
| Naive Prairie | Medial Core | Oxtr Puncta Frequency | Post Hoc Fisher's Exact Test with Bonferonni correction (0.0083) | No Drd vs D1+D2 | na | | 0.1561 | ns | | 3H (left) | R studio |
| | | | | Drd1 vs Drd2 | na | | 0.1719 | ns | | | |
| | | | | Drd1 vs D1+D2 | na | | 0.0009 | ** | | | |
| | | | | Drd2 vs D1+D2 | na | | 0.4299 | ns | | | |
| Naive Prairie | Medial Shell | Oxtr Puncta Frequency | Chi Square Test of Independence | No Drd vs Drd 1 vs Drd2 vs Drd1+Drd2 | 11.2602 | | 0.0104 | ** | | SF 1C (left) | Python |
| | | | Post Hoc Chi-Square Test for Trend with Bonferonni correction (0.0083) | Drd1 vs No Drd | 4.7125 | | 0.1797 | ns | | | |
| | | | | Drd1 vs Drd2 | 2.3127 | | 0.7699 | ns | | | |
| | | | | Drd2 vs No Drd | 7.5125 | | 0.0368 | ns | | | |
| | | | | D1+D2 vs No Drd | 11.1002 | | 0.0052 | * | | | |
| | | | | D1+D2 vs Drd1 | 9.9936 | | 0.0094 | ns | | | |
| | | | | D1+D2 vs Drd2 | 3.4148 | | 0.3877 | ns | | | |
| Naive Prairie | Lateral Shell | Oxtr Puncta Frequency | Generalized Fisher's Exact Test | No Drd vs Drd 1 vs Drd2 vs Drd1+Drd2 | na | | 0.0005 | ** | | SF 1G (left) | R studio |
| | | | Post Hoc Fisher's Exact Test with Bonferonni correction (0.0083) | No Drd vs Drd1 | na | | 1.0000 | ns | | | |
| | | | | No Drd vs Drd2 | na | | 0.6141 | ns | | | |
| | | | | No Drd vs D1+D2 | na | | 0.0818 | ns | | | |
| | | | | Drd1 vs Drd2 | na | | 0.6525 | ns | | | |
| | | | | Drd1 vs D1+D2 | na | | 0.0060 | ** | | | |
| | | | | Drd2 vs D1+D2 | na | | 0.6019 | ns | | | |
| Pairbonded Prairie | Medial Core | Oxtr Puncta Frequency | Chi Square Test of Independence | No Drd vs Drd 1 vs Drd2 vs Drd1+Drd2 | 28.9829 | | <0.0001 | **** | | 3H (right) | Python |
| | | | Post Hoc Chi-Square Test for Trend with Bonferonni correction (0.0083) | Drd1 vs No Drd | 6.0215 | | 0.0848 | ns | | | |
| | | | | Drd1 vs Drd2 | 3.0198 | | 0.4935 | ns | | | |
| | | | | Drd2 vs No Drd | 9.6081 | | 0.0116 | ns | | | |
| | | | | D1+D2 vs No Drd | 20.1709 | | <0.0001 | **** | | | |
| | | | | D1+D2 vs Drd1 | 30.3492 | | <0.0001 | **** | | | |
| | | | | D1+D2 vs Drd2 | 18.2546 | | 0.0001 | *** | | | |
| Pairbonded Prairie | Medial Shell | Oxtr Puncta Frequency | Chi Square Test of Independence | No Drd vs Drd 1 vs Drd2 vs Drd1+Drd2 | 31.0097 | | <0.0001 | **** | | SF 1C (right) | Python |
| | | | Post Hoc Chi-Square Test for Trend with Bonferonni correction (0.0083) | Drd1 vs No Drd | 2.1098 | | 0.8781 | ns | | | |
| | | | | Drd1 vs Drd2 | 13.4526 | | 0.0015 | *** | | | |
| | | | | Drd2 vs No Drd | 8.9579 | | 0.0166 | ns | | | |
| | | | | D1+D2 vs No Drd | 17.8591 | | 0.0001 | **** | | | |
| | | | | D1+D2 vs Drd1 | 43.3264 | | <0.0001 | **** | | | |
| | | | | D1+D2 vs Drd2 | 8.9685 | | 0.0165 | ns | | | |
| Pairbonded Prairie | Lateral Shell | Oxtr Puncta Frequency | Chi Square Test of Independence | No Drd vs Drd 1 vs Drd2 vs Drd1+Drd2 | 47.8407 | | < 0.0001 | **** | | SF 1G (right) | Python |
| | | | Post Hoc Chi-Square Test for Trend with Bonferonni correction (0.0083) | Drd1 vs No Drd | 2.517 | | 0.6528 | ns | | | |
| | | | | Drd1 vs Drd2 | 16.2552 | | 0.0003 | *** | | | |
| | | | | Drd2 vs No Drd | 11.7916 | | 0.0036 | ** | | | |
| | | | | D1+D2 vs No Drd | 26.9644 | | < 0.0001 | **** | | | |
| | | | | D1+D2 vs Drd1 | 64.3111 | | < 0.0001 | **** | | | |
| | | | | D1+D2 vs Drd2 | 16.7138 | | 0.0003 | *** | | | |
| Naive Meadow | Medial Core | Oxtr Puncta Frequency | Generalized Fisher's Exact Test | No Drd vs Drd 1 vs Drd2 vs Drd1+Drd2 | na | | 0.0951 | ns | | 3I (left) | R studio |
| Naive Meadow | Medial Shell | Oxtr Puncta Frequency | Generalized Fisher's Exact Test | No Drd vs Drd 1 vs Drd2 vs Drd1+Drd2 | na | | 0.0006 | *** | | SF 1D (left) | R studio |
| | | | Post Hoc Fisher's Exact Test with Bonferonni correction (0.0083) | No Drd vs Drd1 | na | | 1.0000 | ns | | | |
| | | | | No Drd vs Drd2 | na | | 1.0000 | ns | | | |
| | | | | No Drd vs D1+D2 | na | | 1.0000 | ns | | | |
| | | | | Drd1 vs Drd2 | na | | 1.0000 | ns | | | |
| | | | | Drd1 vs D1+D2 | na | | 0.0080 | ** | | | |
| | | | | Drd2 vs D1+D2 | na | | 0.0035 | ** | | | |
| Naive Meadow | Lateral Shell | Oxtr Puncta Frequency | Generalized Fisher's Exact Test | No Drd vs Drd 1 vs Drd2 vs Drd1+Drd2 | na | | 0.5352 | ns | | SF 1H (left) | R studio |
| Paired Meadow | Medial Core | Oxtr Puncta Frequency | Generalized Fisher's Exact Test | No Drd vs Drd 1 vs Drd2 vs Drd1+Drd2 | na | | 0.1938 | ns | | 3I (right) | R studio |
| Paired Meadow | Medial Shell | Oxtr Puncta Frequency | Generalized Fisher's Exact Test | No Drd vs Drd 1 vs Drd2 vs Drd1+Drd2 | na | | 0.5133 | ns | | SF 1D (right) | R studio |
| Paired Meadow | Lateral Shell | Oxtr Puncta Frequency | Generalized Fisher's Exact Test | No Drd vs Drd 1 vs Drd2 vs Drd1+Drd2 | na | | 0.8230 | ns | | SF 1H (right) | R studio |
| | | | | NT Naive vs NT Pairbonded | 3.4716 | | 0.0061 | ** | 0.4412, 0.8476 | | |

| | | | | | | | | | | | |
|---|---|---|---|---|---|---|---|---|---|---|---|
| Naïve vs Pairbonded Prairie | Medial Core | % Positive Cells, all 8 transcript combinations | t-test on a ratio of means with error propagation & FDR corrected p-value | Drd1 only Naive vs Drd1 only Pairbonded | -1.6152 | | 0.3281 | ns | 0.2790, 1.0739 | SF 2A | Python |
| | | | | Drd2 only Naive vs Drd2 only Pairbonded | 0.5866 | | 0.6386 | ns | 0.6998, 1.5521 | | |
| | | | | Oxtr only Naive vs Oxtr only Pairbonded | 1.5553 | | 0.3281 | ns | 0.6854, 3.5974 | | |
| | | | | D1/D2 Naive vs D1/D2 Pairbonded | 0.0777 | | 0.9382 | ns | 0.3479, 1.7053 | | |
| | | | | D1/Oxtr Naive vs D1/Oxtr Pairbonded | 1.0133 | | 0.5515 | ns | 0.4900, 2.5747 | | |
| | | | | D2/Oxtr Naive vs D2/Oxtr Pairbonded | 0.9495 | | 0.5515 | ns | 0.4419, 2.5825 | | |
| | | | | D1/D2/Oxtr Naive vs D1/D2/Oxtr Pairbonded | 0.7296 | | 0.6231 | ns | 0.3217, 2.4673 | | |
| Naïve vs Pairbonded Prairie | Medial Shell | % Positive Cells, all 8 transcript combinations | t-test on a ratio of means with error propagation & FDR corrected p-value | NT Naive vs NT Pairbonded | 1.9366 | | 0.2225 | ns | 0.9872, 2.0578 | SF 2B | Python |
| | | | | Drd1 only Naive vs Drd1 only Pairbonded | -0.1313 | | 0.8958 | ns | 0.4827, 1.4530 | | |
| | | | | Drd2 only Naive vs Drd2 only Pairbonded | -0.1670 | | 0.8958 | ns | 0.3955, 1.5107 | | |
| | | | | Oxtr only Naive vs Oxtr only Pairbonded | 0.8821 | | 0.6686 | ns | 0.4646, 2.3925 | | |
| | | | | D1/D2 Naive vs D1/D2 Pairbonded | -4.0282 | | 0.0009 | ** | 0.2349, 0.7398 | | |
| | | | | D1/Oxtr Naive vs D1/Oxtr Pairbonded | 0.8135 | | 0.6686 | ns | 0.3606, 2.5283 | | |
| | | | | D2/Oxtr Naive vs D2/Oxtr Pairbonded | -0.5290 | | 0.7973 | ns | 0.2613, 1.4277 | | |
| | | | | D1/D2/Oxtr Naive vs D1/D2/Oxtr Pairbonded | -0.9005 | | 0.6686 | ns | 0.3048, 1.2611 | | |
| Naïve vs Pairbonded Prairie | Lateral Shell | % Positive Cells, all 8 transcript combinations | t-test on a ratio of means with error propagation & FDR corrected p-value | NT Naive vs NT Pairbonded | -0.5784 | | 0.7441 | ns | 0.5021, 1.273 | SF 2C | Python |
| | | | | Drd1 only Naive vs Drd1 only Pairbonded | -1.5472 | | 0.6945 | ns | 0.2166, 1.0969 | | |
| | | | | Drd2 only Naive vs Drd2 only Pairbonded | -0.3274 | | 0.7441 | ns | 0.5406, 1.3293 | | |
| | | | | Oxtr only Naive vs Oxtr only Pairbonded | 1.2960 | | 0.6945 | ns | 0.1121, 5.2330 | | |
| | | | | D1/D2 Naive vs D1/D2 Pairbonded | -0.3317 | | 0.7441 | ns | 0.3113, 1.4914 | | |
| | | | | D1/Oxtr Naive vs D1/Oxtr Pairbonded | 0.7854 | | 0.6945 | ns | 0.5402, 2.0622 | | |
| | | | | D2/Oxtr Naive vs D2/Oxtr Pairbonded | 0.9309 | | 0.6945 | ns | 0.4176, 2.6121 | | |
| | | | | D1/D2/Oxtr Naive vs D1/D2/Oxtr Pairbonded | 0.8511 | | 0.6945 | ns | 0.3129, 2.7196 | | |
| Naïve vs Paired Meadow | Medial Core | % Positive Cells, all 8 transcript combinations | t-test on a ratio of means with error propagation & FDR corrected p-value | NT Naive vs NT Paired | 2.1784 | | 0.0846 | ns | 1.0514, 2.1015 | SF 2A | Python |
| | | | | Drd1 only Naive vs Drd1 only Paired | 1.3142 | | 0.3836 | ns | 0.8594, 1.6922 | | |
| | | | | Drd2 only Naive vs Drd2 only Paired | -3.8622 | | 0.0016 | ** | 0.6661, 0.8927 | | |
| | | | | Oxtr only Naive vs Oxtr only Paired | 0.8614 | | 0.4469 | ns | 0.2905, 2.7984 | | |
| | | | | D1/D2 Naive vs D1/D2 Paired | -2.2258 | | 0.0846 | ns | 0.3986, 0.9654 | | |
| | | | | D1/Oxtr Naive vs D1/Oxtr Paired | 1.0823 | | 0.4266 | ns | 0.1869, 3.7651 | | |
| | | | | D2/Oxtr Naive vs D2/Oxtr Paired | -0.0652 | | 0.9481 | ns | 0.1055, 1.8375 | | |
| | | | | D1/D2/Oxtr Naive vs D1/D2/Oxtr Paired | -0.9995 | | 0.4266 | ns | 0.0779, 1.3042 | | |
| Naïve vs Paired Meadow | Medial Shell | % Positive Cells, all 8 transcript combinations | t-test on a ratio of means with error propagation & FDR corrected p-value | NT Naive vs NT Paired | 2.5140 | | 0.0541 | | 1.1898, 2.6102 | SF 2B | Python |
| | | | | Drd1 only Naive vs Drd1 only Paired | 1.0898 | | 0.4249 | | 0.8060, 1.6668 | | |
| | | | | Drd2 only Naive vs Drd2 only Paired | 0.1483 | | 0.4249 | | 0.5729, 1.1611 | | |
| | | | | Oxtr only Naive vs Oxtr only Paired | 1.3523 | | 0.3658 | | 0.6864, 2.6560 | | |
| | | | | D1/D2 Naive vs D1/D2 Paired | -3.6401 | | 0.0035 | ** | 0.3991, 0.8230 | | |
| | | | | D1/Oxtr Naive vs D1/Oxtr Paired | 0.9373 | | 0.4249 | | 0.3652, 2.7717 | | |
| | | | | D2/Oxtr Naive vs D2/Oxtr Paired | 0.1803 | | 0.8573 | | 0.3350, 1.7979 | | |
| | | | | D1/D2/Oxtr Naive vs D1/D2/Oxtr Paired | -1.3411 | | 0.3658 | | 0.2424, 1.1465 | | |
| Naïve vs Paired Meadow | Lateral Shell | % Positive Cells, all 8 transcript combinations | t-test on a ratio of means with error propagation & FDR corrected p-value | NT Naive vs NT Paired | 0.8940 | | 0.4980 | | 0.7623, 1.6275 | SF 2C | Python |
| | | | | Drd1 only Naive vs Drd1 only Paired | 3.0992 | | 0.0056 | ** | 1.2652, 2.2089 | | |
| | | | | Drd2 only Naive vs Drd2 only Paired | -3.0634 | | 0.0056 | ** | 0.5932, 0.9130 | | |
| | | | | Oxtr only Naive vs Oxtr only Paired | -0.1601 | | 0.8731 | | 0.0762, 1.7858 | | |
| | | | | D1/D2 Naive vs D1/D2 Paired | -3.0945 | | 0.0056 | ** | 0.4530, 0.8804 | | |
| | | | | D1/Oxtr Naive vs D1/Oxtr Paired | 0.3165 | | 0.8597 | | 0.5106, 1.6752 | | |
| | | | | D2/Oxtr Naive vs D2/Oxtr Paired | -2.8353 | | 0.0089 | ** | 0.0259, 0.8279 | | |
| | | | | D1/D2/Oxtr Naive vs D1/D2/Oxtr Paired | -5.0947 | | 0.0000 | **** | -0.0067, 0.5576 | | |
